# Long-Lasting Electrohydrodynamically Printed Transparent Soft Microelectrode for Implantable Biointerfaces

**DOI:** 10.64898/2026.05.19.726391

**Authors:** Hyeongjin Jo, Gaeun Lee, Yujun Song, Seo Yeon Kim, Minho Kim, Rahul Manna, Dayeong Choi, Abiodun Aderibigbe, Steven L. Suib, Kiyeon Park, Jungho Ahn, Ji-Hyeon Song, Kyungjin Kim

**Author notes:** Equally contributed (Hyeongjin Jo, Gaeun Lee).

## Abstract

Reliable and scalable soft implantable neural interface fabrication remains a key challenge for chronic bioelectronic applications. Here, we present a transparent soft microelectrode fabricated with electrohydrodynamic (EHD) printing, utilizing the fluorinated polymer poly(vinylidene fluoride-co-hexafluoropropylene) (PVDF-HFP) and poly (3, 4-ethylenedioxythiophene) polystyrene sulfonate (PEDOT: PSS) to form seamless, selectively patterned multilayer structures with low impedance and long-term stability. Controlled in situ curing during printing yields dense, void-free substrate and encapsulation layers, suppressing interfacial defects and ionic pathways, while maintaining high optical transparency (>60%) with PEDOT:PSS. The printed microelectrodes exhibit low impedance, high charge storage and injection capacities, and stable electrochemical behavior under biomimetic conditions. In addition, the devices demonstrate robust mechanical and electromechanical stability under cyclic deformation in both dry and wet environments, as well as under prolonged electrical stimulation. Accelerated aging studies project multi-year operational lifetimes, and in vitro/in vivo biocompatibility assessments confirm excellent tissue integration. These results establish EHD-printed fluorinated polymer-based microelectrodes as a scalable and durable platform for chronic implantable biointerfaces.

**ToC:** 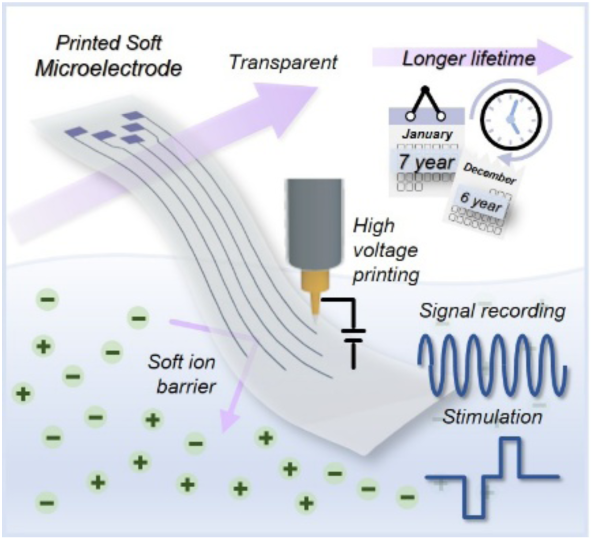

This report presents an electrohydrodynamically printed transparent soft microelectrode for chronic purposes. Electrohydrodynamic printing promotes seamless multilayer structures with selective deposition and long-term mechanical stability. The devices show low impedance, high charge capacity, and robust electrochemical/electromechanical properties. Accelerated aging projects ∼7.2 year lifetimes, and XPS/SEM-EDS confirm strong ion barrier properties and biocompatibility for chronic implantation.

## 1. Introduction

Advances in biointerfaces for diagnostics and human–machine interfaces are paving the way for improved bio-signal recording and interpretation in both clinical and scientific applications.^[1–4]^ In particular, soft and conductive materials capable of long-term operation are increasingly explored for chronic implantable devices.^[5–7]^ Moreover, soft biointerfaces mitigate key limitations of conventional silicon-based stiff and rigid form factors, including foreign-body responses such as scar formation that compromises long-term performance.^[3,4]^

Implantable microelectrodes have traditionally been fabricated using various manufacturing approaches, including lithography^[7–9]^, 3D printing^[10–12]^, and laser processing^[13,14]^. While lithography-based microfabrication enables high spatiotemporal resolution with excellent reproducibility, it involves complex, labor-intensive processes that increase fabrication cost and introduce significant production downtime between processing steps.^[15]^ In contrast to these conventional approaches, printing-based techniques offer simplified processing, enabling on-demand production with facile customization and improved scalability.^[11,12]^ Printing techniques enable rapid, batch fabrication of neural interface devices, mitigating major obstacles of conventional microfabrication processes.^[12,15]^ Among them, electrohydrodynamic (EHD) printing is particularly advantageous for device integration and conformal deposition, as it utilizes an electric field between the nozzle and substrate to support high-resolution microscale patterning beyond typical inkjet printing, offering a pathway toward finer conductive features for future high-density biointerface applications.^[16,17]^ This printing technique further enables robust interlayer integration within multilayer device architectures, which is a critical design factor for implantable electrodes.^[18]^

Metal-free organic transparent microelectrodes offer advantages in terms of post-implantation optical access, in contrast to metal-based electrodes, enabling optical stimulation while maintaining compatibility with medical imaging modalities such as magnetic resonance imaging (MRI) system.^[19,20]^ Furthermore, conventional opaque metal-based electrodes may cause photoelectric artifacts, which lead to electrophysiological signal interference when applying optogenetics to treat neurodegenerative diseases such as Parkinson’s disease and depression.^[21–23]^ As such, the adoption of transparent materials for neural interfaces to fabricate fully transparent implants is necessary for advanced biomedical systems. In line with this trend, PEDOT:PSS has been widely employed as a conductive material due to its high electrochemical stability and corrosion resistance in aqueous environments, while also offering optical transparency for optical stimulation and post-implantation tissue inspection.^[18,24–26]^ For encapsulation material, fluorinated polymers are emerging as strong candidates as they effectively suppress ion penetration in ionic environments. Owing to this characteristic, fluorinated polymers have been successfully applied in bioelectronics and energy storage systems requiring stability in ionic environments.^[7,27]^ Among these, polyvinylidene fluoride (PVDF) and its copolymer poly(vinylidene fluoride-co-hexafluoropropylene) (PVDF-HFP) have emerged as promising materials, as their strong C-F bonding imparts mechanical robustness, chemical stability, and resistance to harsh environments. ^[28–30]^ However, combining two different materials into a single system using additive methods is challenging, particularly when maintaining structural integrity due to differences in chemical composition that lead to variations in surface characteristics.^[31–33]^ To mitigate these issues, mechanical interlocking methods or surface treatments are commonly employed.^[32–35]^ However, these approaches may increase the complexity of the manufacturing process or compromise the inherent advantages of the materials. Therefore, integrating these material advantages with simplified scalable manufacturing while maintaining interfacial integrity and long-term stability remains a key challenge.

Herein, we introduce a transparent and soft microelectrode fabricated via EHD printing, utilizing the fluorinated polymer PVDF-HFP and the conductive polymer PEDOT:PSS to form seamless, selectively patterned multilayer structures with low impedance and long-term stability. A controlled curing strategy during printing enables dense, void-free substrate and encapsulation layers, minimizing interfacial defects and suppressing ionic transport pathways, while supporting conformal PEDOT:PSS deposition and maintaining high optical transparency (>60%). To assess device performance, electrochemical, mechanical, and electromechanical characterizations were conducted. The printed microelectrodes exhibit low impedance, high charge storage and injection capacities, and stable electrochemical performance under biomimetic conditions. Accelerated aging studies reveal projected multi-year operational lifetimes (∼7.2 years), exceeding previously reported PEDOT:PSS-based soft microelectrodes. Additionally, the compatibility of lifetime testing approaches under elevated temperatures for polymer-based microelectrodes was assessed. In vitro and in vivo biocompatibility evaluations further confirm excellent cell growth as well as tissue integration, establishing a scalable and durable platform for chronic implantable biointerfaces.

## 2. Result and discussion

### 2.1. Electrohydrodynamically printed transparent soft microelectrode

The schematic illustration of the EHD printing process for the transparent soft microelectrode is shown in **Figure 1a**, **Figure S1**, and **Video S1** (Supporting Information). The printing process began with preheating the heat bed to 40 °C to enable controlled in-situ crystallization of PVDF-HFP and gradual solvent evaporation. This controlled curing suppresses excessive ink spreading on the sacrificial polyethylene terephthalate (PET) substrate and promotes seamless integration between consecutively printed layers. In contrast, excessively high heat-bed temperatures accelerate solvent evaporation, inducing porous structures in the PVDF-HFP film that can lead to ionic penetration under implanted conditions, rendering it unsuitable as an ionic barrier. Additionally, elevated temperatures can cause premature solidification of the ink near the nozzle outlet, causing clogging, which is undesirable for reliable batch production.

**Figure 1.**
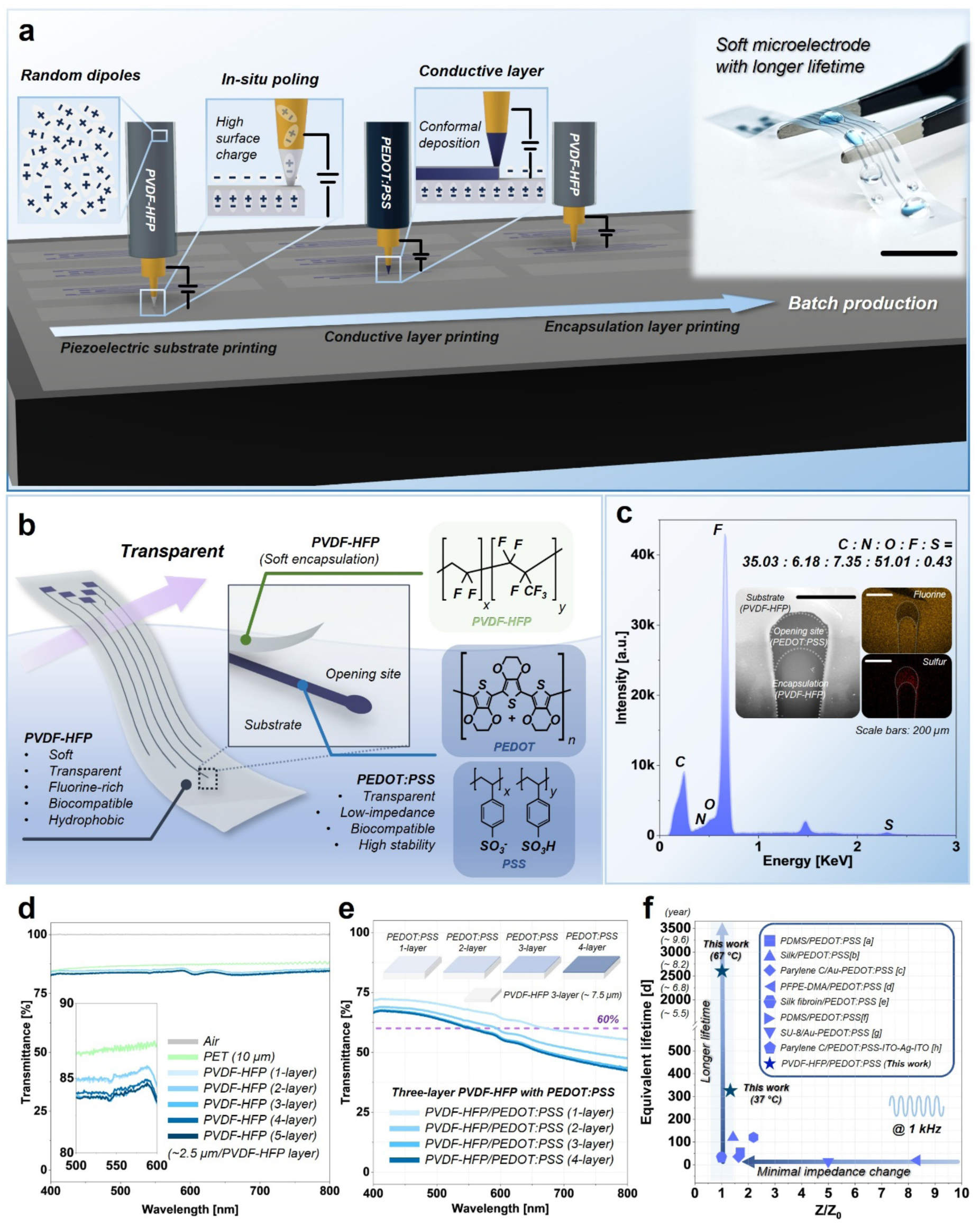
a) Schematic illustration of electrohydrodynamic printing process of soft microelectrodes with batch production. Inset: photograph image of a printed microelectrode. Scale bar: 1 cm. b) Overview of the electrohydrodynamically printed soft microelectrode with an exploded view near opening site with chemical structure of materials. c) SEM–EDS analysis of the substrate and opening sites. Inset: SEM image and elemental mapping results. d) Air-based transmission spectra of PET film and PVDF-HFP films with different numbers of printed layers. Inset: magnified transmittance near 550 nm. e) Transmittance spectra of tri-layer PVDF-HFP film with different PEDOT:PSS printed layers. f) Comparison of impedance changes over time and corresponding lifetimes at 1 kHz with previously reported PEDOT:PSS-based soft microelectrodes. [a] ^[36]^, [b] ^[37]^, [c]^[38]^, [d]^[39]^, [e]^[40]^, [f]^[41]^, [g]^[42]^, [h]^[43]^.

By employing a slow, partial curing strategy, the substrate thickness can be controlled by adjusting the number of printing cycles. As shown in **Figure S2**, the final printed substrate exhibits seamless layer integration, enhancing mechanical robustness while minimizing interlayer delamination. Cross-sectional SEM images further show a dense, void-free structure after final curing, as well as high integration among the layers (**Figure S2**). In addition, the high-voltage EHD printing process enables in-situ poling, as demonstrated in our previous study^[17]^, which enhances surface charge and facilitates conformal deposition of the conductive PEDOT:PSS layer, ensuring stable device operation. The following printing step involved selective deposition of an encapsulation layer on top of the conductive traces, improving material efficiency and reducing processing steps. The final curing step at elevated temperature facilitates complete solvent evaporation and further polymer crosslinking, enhancing structural integrity. The advantages of using PVDF-HFP and PEDOT:PSS lie in their complementary material properties under ionic environments. PVDF-HFP acts as a robust encapsulation barrier due to its fluorinated, hydrophobic nature, delaying water sorption, while PEDOT:PSS is hydrophilic, enabling effective interfacial contact with biological tissues (**Figure 1b, Figure S3**).

As shown in **Figure 1b** and **Figure S4-S5**, the printed device consists of five channels and three layers: substrate, conductive, and encapsulation layers. The size of the opening site was measured as approximately 115,700 μm^2^ (**Figure S4**). **Figure 1c** presents the surface SEM– EDS results near the opening site. The opening region exhibits sulfur-rich characteristics originating from PEDOT:PSS, while both the substrate and encapsulation layers show strong fluorine signals corresponding to PVDF-HFP. **Figure 1d** shows the optical transmission spectra of PVDF-HFP films with different printed layers, compared with a PET film, which is commonly used as a transparent reference substrate. The transmittance values of PVDF-HFP films with one to five layers were measured as 85.1, 85.0, 83.8, 84.1, and 84.0 % at a wavelength of 550 nm under air, respectively, indicating no significant dependence on the number of printed layers. The PET film exhibited a transmittance of 86 %, demonstrating that the transparency of the PVDF-HFP film is comparable. This consistently high transmittance suggests a dense and uniform film morphology, minimizing light scattering typically associated with porous structures.

Further transmittance analysis was conducted after depositing the PEDOT:PSS layer on top of the PVDF-HFP tri-layer to evaluate interlayer integrity and optical transparency. **Figure 1e** presents the transmittance spectra of multilayer films with varying PEDOT:PSS printing cycles. As the number of PEDOT:PSS printing cycles increased, the transmittance at 550 nm decreased to 67.8%, 63.1%, 60.7%, and 60.3%, respectively. As expected, increasing the PEDOT:PSS layer thickness reduced the optical transparency. Nevertheless, devices printed with up to quad PEDOT:PSS layers maintained transmittance values above 60%, indicating excellent optical transparency even in multilayer configurations (**Figure 1e**). Moreover, the transmittance spectra exhibited no noticeable light scattering or haze, demonstrating conformal deposition and high interfacial integrity without defect-induced optical loss. The robustness of the EHD-printed microelectrodes under in vitro lifetime testing is further supported by comparing with previously reported PEDOT:PSS-based microelectrodes.^[36–43]^ **Figure 1f** presents the equivalent aging time as a function of impedance variation for the PEDOT:PSS-based microelectrodes developed in this work and previous reports.^[36–43]^ The electrohydrodynamically printed microelectrodes demonstrate superior stability, exhibiting consistent impedance values under both in vitro 1× PBS testing at body temperature and accelerated aging conditions, compared to previously reported soft PEDOT:PSS microelectrodes.^[36–43]^ A detailed discussion of long-term stability and failure mechanisms is provided in **Section 2.4**.

### 2.2. Electrochemical performance and durability of electrohydrodynamically printed soft microelectrode

The electrochemical performance of the implantable microelectrode is paramount, as it reflects the capability of the device for signal recording as well as stimulation of the biological system. The basic electrochemical performance of the EHD-printed device was evaluated using electrochemical impedance spectroscopy (EIS), cyclic voltammetry (CV), and voltage transient (VT) measurements. For the initial characterization stage, the electrochemical properties were measured in 1× PBS at 37 °C to simulate physiological conditions. **Figure 2a** demonstrates the EIS and CV measurement conditions, which were performed using a three-electrode setup including Ag/AgCl, Pt, and the printed device as the reference, counter, and working electrode, respectively, under controlled temperature conditions.

**Figure 2.**
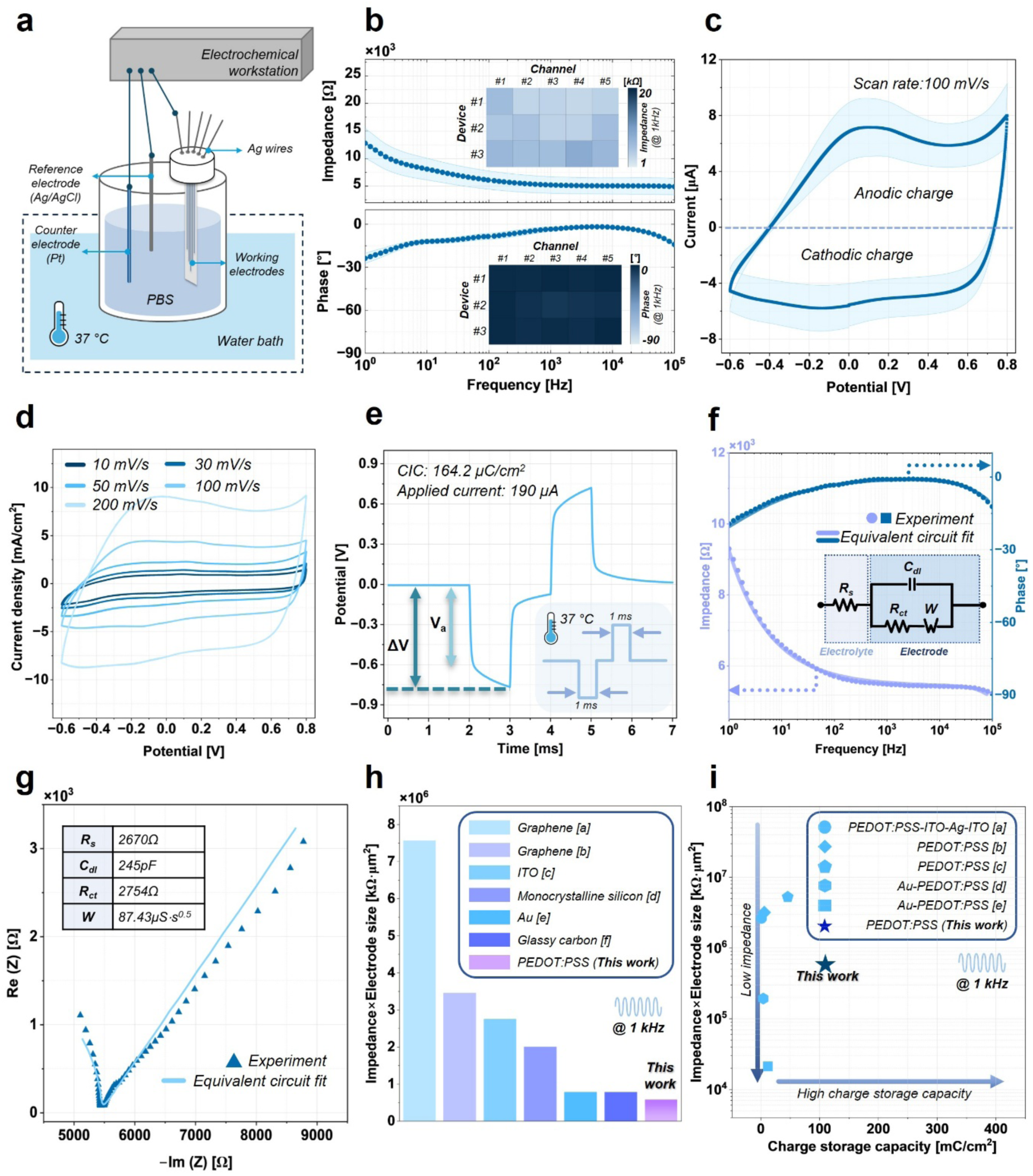
a) Schematic illustration of the electrochemical setup. b) Bode plot of the device. c) Cyclic voltammetry measurement at a scan rate of 100 mV s⁻¹. The shaded area represents the standard deviation (n = 15 electrodes). d) Current density measured at various scan rates. e) Voltage transient measurement. Equivalent circuit fitting of experimental data: f) Impedance and phase, and g) Nyquist plot. Inset: equivalent circuit. All tests were conducted in 1× PBS at 37 °C unless otherwise stated. h, i) Electrochemical property comparison with previous works: h) Normalized impedance at 1 kHz among various conductive materials; [a] ^[44]^, [b] ^[45]^, [c] ^[46]^, [d] ^[47]^, [e] ^[48]^, [f] ^[49]^. i) Normalized impedance at 1 kHz and charge storage capacity of previously reported PEDOT:PSS-based microelectrodes; [a] ^[50]^, [b] ^[51]^, [c] ^[52]^, [d] ^[53]^, [e] ^[54]^.

**Figure 2b** shows the Bode plot recorded through EIS measurements with 15 electrodes over the frequency range from 10^0^ to 10^5^ Hz. The impedance values at 10 Hz, and 1 kHz and 100 kHz were measured as 8.1 ± 1.9, 5.2 ± 1.5, and 4.9 ± 1.5 kΩ, respectively, indicating low impedance across a broad range of frequency range relevant to neural interfacing. The CV measurements exhibit a near-square shape with a stable response within the safe voltage window of –0.6 to 0.8 V, with a calculated charge storage capacity (CSC) of 110 ± 11.7 mC/cm^2^ (**Figure 2c**). The near rectangular CV profile suggests capacitive charge transfer behavior and stable electrode–electrolyte interaction. Additional CV measurements at different scan rates show consistent curve shapes without significant distortion, confirming electrochemical stability (**Figure 2d**). Diffusion-controlled behavior was further validated by plotting anodic and cathodic peak currents against the square root of the scan rate, showing a linear relationship (**Figure S6**).

Long-term electrochemical stability was assessed through repeated CV cycling, where the device was subjected to 1,400 cycles (**Figure S7**). The results showed minimal capacitance degradation, with approximately 93% of the initial value retained and no significant increase in impedance, corresponding to a normalized impedance (Z/Z_0_) of 1.11 at 1 kHz. These results indicate stable electrochemical performance under repeated operation.

VT measurements were further conducted to calculate the charge injection capacity (CIC), which is critical for stimulation applications (**Figure 2e**). The applied current was adjusted to remain within the safe voltage limits utilized for CV measurement. The device exhibited rapid and stable voltage responses with near-square waveforms, indicating efficient charge injection behavior. The CIC value was calculated as 162 μC/cm², demonstrating a high CIC performance (**Figure 2e**). Additionally, cyclic VT stimulation testing over 10,000 cycles showed consistent waveform profiles without degradation, confirming stable stimulation performance (**Figure S8**).

Another important aspect of stability in aqueous environments is the sorption behavior of the device. When operated in biofluid environments, swelling can lead to device drift and interfacial delamination, ultimately degrading signal reliability. In addition, water uptake facilitates ion transport, potentially leading to electrical failure. To evaluate these effects, sorption testing was performed by measuring weight gain over time in deionized (DI) water. As shown in **Figure S9**, the device exhibited minimal weight gain of approximately 0.2 mg (∼2 %), reaching saturation within ∼40 minutes, indicating limited water uptake and stable structural integrity. The primary weight increase is attributed to water absorption through the hydrophilic PEDOT:PSS opening sites, with a minor contribution from surface adsorption on the hydrophobic PVDF-HFP.

To further understand the electrochemical behavior, an equivalent circuit model based on the Randles circuit was constructed (**Figure 2f–g**). The model includes solution resistance (R_s_), double-layer capacitance (C_dl_), charge-transfer resistance (R_ct_), and Warburg impedance (W). The fitted Bode and Nyquist plots showed good agreement with experimental data, validating the suitability of the model for describing the device behavior in PBS.

Impedance is a critical parameter for implantable electrodes, as it directly influences signal quality during recording. **Figure 2h** compares the normalized impedance at 1 kHz of commonly used electrode materials with that of the EHD-printed PEDOT:PSS microelectrode.^[44–49]^ The device presented in this work exhibits comparatively lower normalized impedance. Furthermore, comparison with previously reported PEDOT:PSS-based microelectrodes, based on both normalized impedance and CSC, demonstrates that the device presented in this work achieves competitive electrochemical performance relative to prior reports (**Figure 2i**).^[50–54]^

### 2.3. Mechanical/electromechanical performances and mechanical durability of printed soft microelectrode

For the next characterization of the printed device, the evaluation involved assessing its mechanical and electromechanical properties, as well as its stability under continuous mechanical deformation in moisture-rich environments. Two types of testing were conducted: dry and wet conditions. For dry testing, the device was evaluated under ambient air without moisture exposure, whereas wet testing was conducted in 1× PBS at 37 °C. The mechanical robustness of the device was first assessed through monotonic tensile testing under dry conditions. **Figure 3a** shows the representative stress–strain curve of the device under applied strain. The tensile test was performed until device failure while both ends were fixed to the jig mounted on the micro-tensile testing machine. As shown in **Figure 3a**, the stress–strain response exhibited an initial linear elastic region up to ∼5% strain, followed by a plateau region up to ∼10% strain, indicating the onset of plastic deformation, after which the device failed at approximately 35% strain. Based on these results, the Young’s modulus, tensile strength, elongation at break, and toughness were calculated as 532.8 ± 93.6 MPa, 15.7 ± 1.7 MPa, 33.8 ± 3.8%, and 4.7 ± 1 MJ/m³, respectively (**Figure 3b**).

**Figure 3.**
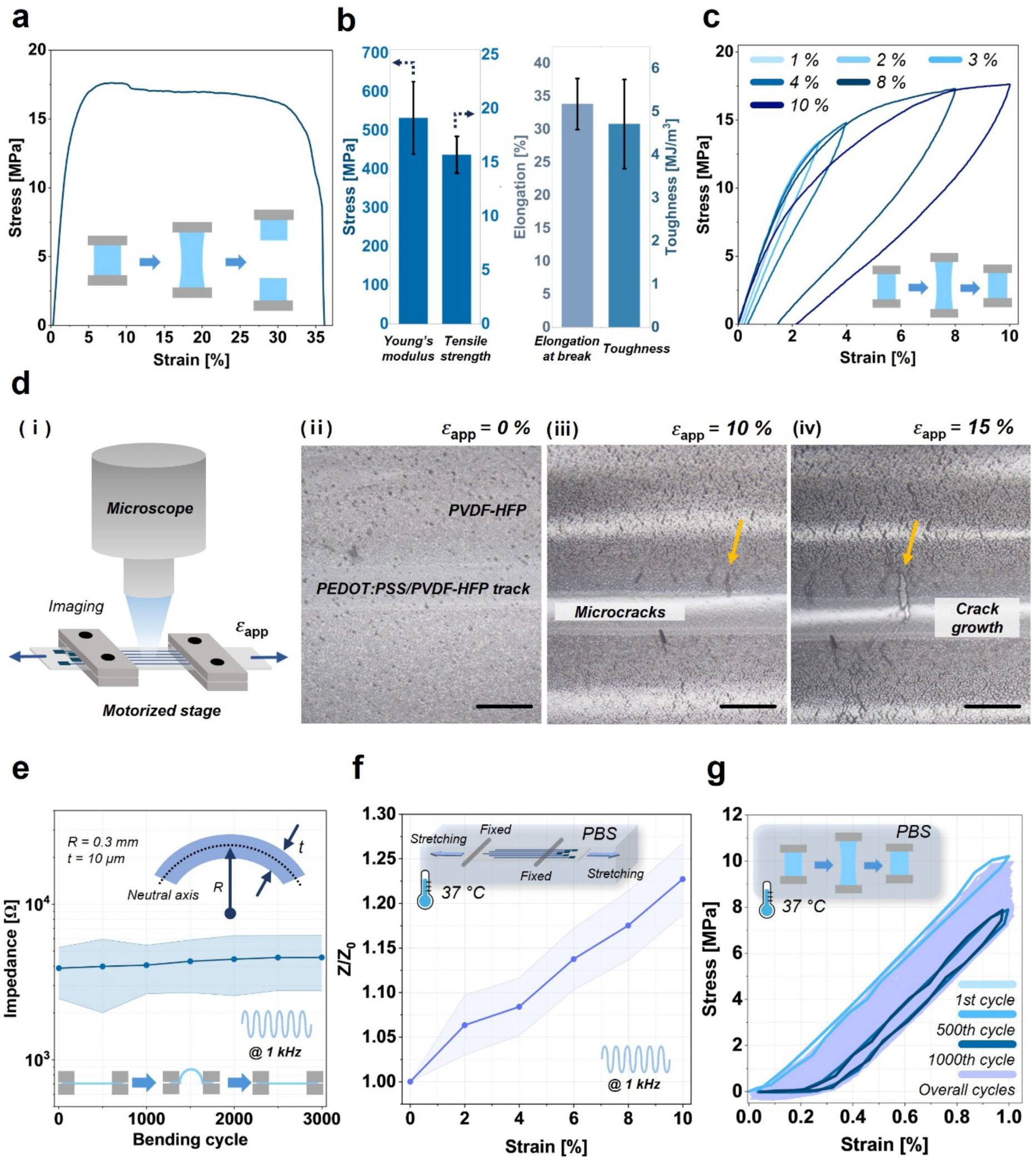
a) Representative stress–strain curve from monotonic tensile testing. Inset: schematic of the testing sample. b) Calculated mechanical properties from tensile testing: Young’s modulus, tensile strength, elongation at break, and toughness. Error bars represent standard deviation (n = 3). c) Cyclic loading test results under various applied strains (1, 2, 3, 4, 8, and 10%). d) Crack onset testing: (i) schematic of the testing setup and microscopic images of (ii) the initial state, applied strain of (iii) 10%, and (iv) 15%. Scale bars: 300 µm. e) Impedance change under mechanical bending (3000 cycles). Inset: schematic of the sample during cyclic bending testing. f) Impedance change under applied strain in 1× PBS at 37 °C. Inset: schematic of the testing setup. Shaded area represents standard deviation (n = 5). g) Stress–strain curve from cyclic mechanical testing in 1× PBS at 37 °C (1000 cycles). Inset: schematic illustration of testing sample under cyclic loading.

Further analysis of the mechanical performance included cyclic loading to determine the strain threshold associated with permanent deformation. Based on the monotonic tensile test results, cyclic loading was applied in the range of 1% to 10% strain (**Figure 3c**). As expected, permanent deformation occurred when strain exceeded ∼8%, as evidenced by residual strain after unloading. The morphological evolution of the device under strain was analyzed using in situ optical microscopy synchronized with the tensile testing system (**Video S2**). **Figure 3d (i)** illustrates the testing setup, while microscopic observations show that initial crack formation in the conductive track and encapsulation layer begins at ∼10% strain (**Figure 3d (ii)**), followed by progressive crack propagation leading to failure (**Figure 3d (iii)**, **Video S2**).

As the final dry characterization of device, cyclic bending tests were performed to assess device stability under continuous mechanical deformation. The device was bent to a radius of curvature of 0.3 mm, corresponding to 1% bending strain, and impedance at 1 kHz was recorded every 500 cycles up to 3000 cycles (**Figure 3e**). The impedance values before and after testing showed no significant change, with values of 3.8 ± 1.4 and 4.5 ± 1.7 kΩ (Z/Z_0_ = 1.18), respectively, indicating stable performance under continuous mechanical deformation and no damage to the conductive trace (**Figure 3e**, **Figure S10**). This result highlights the mechanical robustness of the soft encapsulation; the absence of a significant decrease in impedance suggests that there were no delamination, cracking, or other structural defects, and that the multilayer device structure remained intact and prevented ion penetration.

To further evaluate electromechanical robustness, mechanical deformation was applied under 1× PBS at 37 °C. **Figure 3f** shows the schematic of the static mechanical testing setup. The device was stretched with manual linear stage from 0% to 10% strain in 2% increments, and the corresponding impedance values were collected (**Figure 3f**, **Figure S11**). The results show a moderate impedance increase at 10% strain, with a normalized value (Z/Z₀) of 1.22 ± 0.03 at 1kHz (n=5). Despite the onset of microcrack formation near this strain level observed in dry testing, the electrochemical performance remained relatively stable under wet conditions, indicating functional tolerance to moderate deformation.

Dynamic mechanical stability was further evaluated through cyclic testing under 1× PBS while simultaneously monitoring mechanical response and impedance (**Figure 3g, Video S3**). The stress–strain response over 1000 cycles shows slight permanent deformation during initial cycles, after which the mechanical response stabilizes. The maximum stress decreased from 10.2 MPa at the first cycle to 8 MPa after ∼50 cycles and remained stable at 7.8 MPa at the 1000th cycle. This stabilization behavior suggests initial structural rearrangement of the polymer under cyclic loading, followed by a stable mechanical response. Testing was conducted at 37 °C using a temperature-controlled reservoir filled with PBS, with the experimental setup and corresponding impedance results shown in **Figure S12**. Impedance measurements before and after cyclic loading show slight increases across frequencies. The average impedance values before and after testing were 7.4 ± 1.3 and 8.9 ± 0.9 kΩ at 10 Hz, 5.6 ± 1.2 and 7.2 ± 0.38 kΩ at 1 kHz, and 4.9 ± 1.0 and 5.8 ± 0.5 kΩ at 100 kHz. The corresponding average normalized impedance values (Z/Z₀) at 10 Hz, 1 kHz, and 100 kHz were 1.20, 1.28, and 1.18, respectively. In all groups, no statistically significant differences were observed across all frequency ranges, suggesting no critical degradation of device performance under continuous mechanical stimulation and compatibility with dynamic implanted environments.

### 2.4. In vitro lifetime analysis of printed soft microelectrodes and compatibility of high-temperature accelerated aging tests for organic polymer–based microelectrodes

For an implantable device intended for chronic use, operational lifetime is critical to ensuring reliable performance after implantation. Furthermore, identifying the failure modes that contribute to long-term loss of device functionality is essential, as this information provides a roadmap for future design improvements. In this study, the lifetime of printed soft microelectrodes was evaluated by fully soaking the devices in 1× PBS and monitoring impedance and capacitance over time. For this testing, sample vials were prepared by filling them with 1× PBS, and the connected silver wires were passed through holes in the caps for data collection (**Figure S13**). The holes were further sealed with adhesive and silicone to secure the wires and prevent PBS evaporation. For the device storage and testing, due to the highly hydrophobic nature of the device, a metal clip was attached to the edge of the substrate to ensure full immersion to PBS; otherwise, the device floated on the PBS surface, leading to inaccurate lifetime evaluation due to incomplete electrolyte exposure (**Figure S13**).

**Figure 4a** summarizes representative failure modes of the printed microelectrodes. The possible failure modes of the devices include, but are not limited to, conductivity loss of PEDOT:PSS, opening site damage, wiring failure, and encapsulation delamination, cracking, and degradation, which allow ion penetration and can lead to short circuits (**Figure 4a**). The assessment of device longevity under implantation conditions began with analysis of impedance trends over time by soaking the devices at three different temperatures: 37, 67, and 87 °C. The 37 °C condition was used as the baseline, corresponding to physiological body temperature, while the other two temperatures represent accelerated aging conditions that increase chemical reaction rates at elevated temperatures. The thermal aging evaluation is based on the assumption that material degradation follows the Arrhenius reaction model, described as ^[55]^:

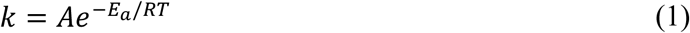

where k, A, Ea, R, and T represent the reaction rate constant, pre-exponential factor, activation energy, gas constant, and absolute temperature (in Kelvin), respectively. Using the Arrhenius relation, the aging acceleration can also be expressed using the Q_10_ factor, where the acceleration factor (AF) is determined by raising Q_10_ to the power of the temperature difference between the testing temperature (T) and physiological temperature (37 °C)^[55]^.

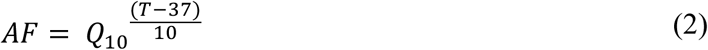

Here, Q_10_ = 2 is commonly used^[56]^ to estimate the equivalent 37 °C lifetime of devices tested under accelerated aging, and this value was applied to estimate the equivalent lifetime of the device in this study. This approach assumes that degradation mechanisms remain consistent across temperatures, which may not strictly hold for polymer-based systems at elevated temperatures.

**Figure 4b** and **Figure S14** show impedance trends under three temperature conditions at three frequencies: 10 Hz, 1 kHz, and 100 kHz. Here, the device failure point was defined as a normalized impedance change (Z/Z₀) exceeding 25, corresponding to an impedance of ∼125 kΩ, which remains within the range of impedance values for “functional” electrodes.^[18]^ As expected, the earliest failures occurred in the 87 °C group, characterized primarily by a rapid and abrupt increase in impedance rather than gradual degradation, across all three frequencies (**Figure 4b** and **Figure S14**). In contrast, devices in the lower-temperature groups (37 and 67 °C) remained stable, with impedance values within the acceptable operational range over time, except for channel #4 in the 67 °C group, for which data could not be collected after Day 292 due to wire detachment and loss from the vial.

**Figure 4.**
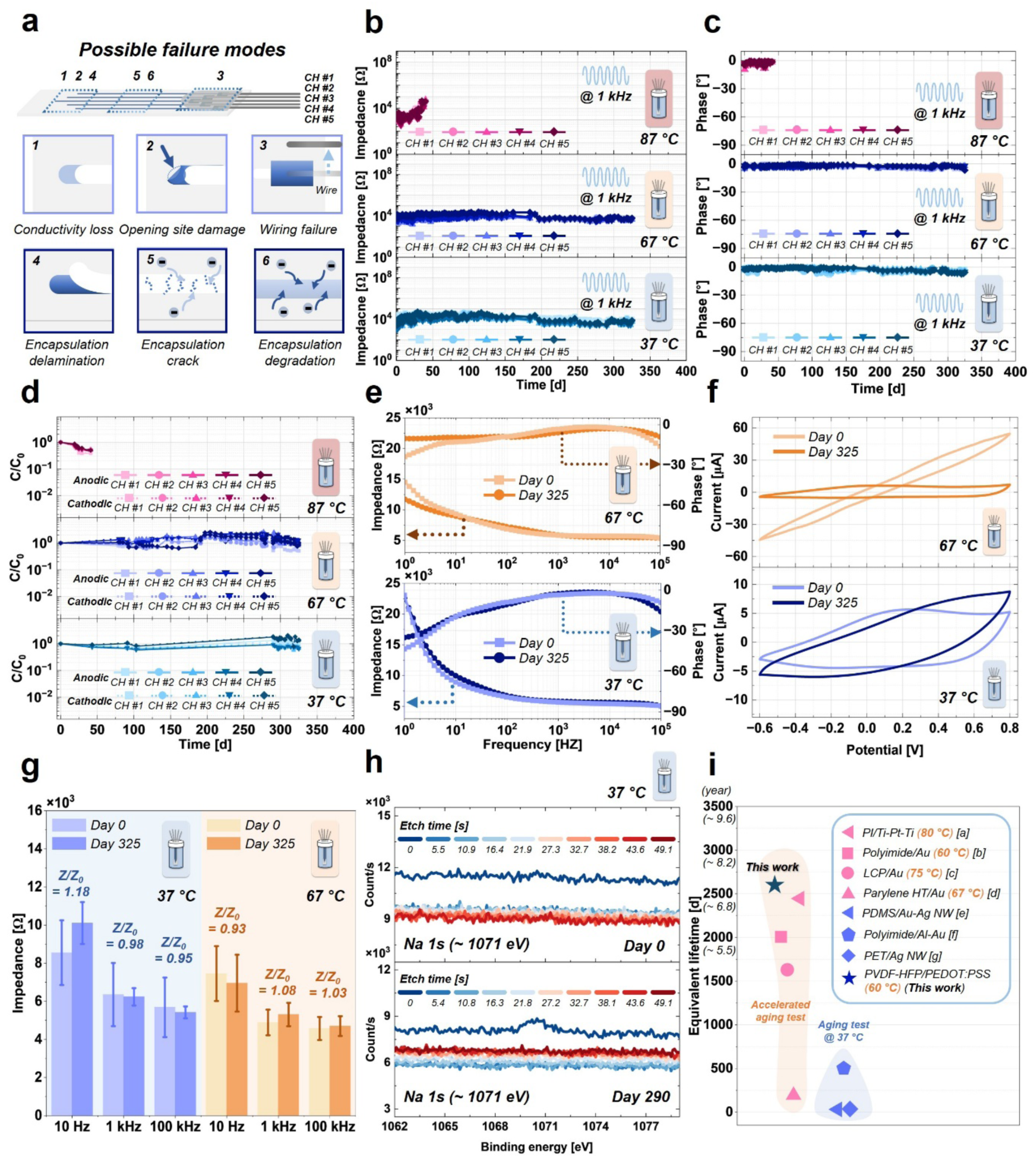
In vitro lifetime testing results. a) Schematic illustration and summary of possible device failure modes. b) Impedance and c) phase change over time in 1X PBS at 1 kHz for three temperature groups: 37, 67, and 87 °C. d) Capacitance change over time for the three temperature groups. Representative e) Bode plots and f) CV curves of post-fabrication and after 325 days of soaking in 1X PBS at 37 (bottom) and 67 °C (top). g) Average impedance at 10 Hz, 1 and 100 kHz before and after 325 days of soaking in 1X PBS at 37 (left) and 67 °C (right). Error bars represent the standard deviation (n = 4). h) XPS spectra for Na 1s before (top) and after soaking (bottom) in 1X PBS at 37 °C for 290 days. i) Comparison of device lifetime with previously reported implantable microelectrodes (format: substrate/conductive material); [a] ^[59]^, [b] ^[60]^, [c] ^[61]^, [d] ^[62]^, [e] ^[63]^, [f] ^[64]^, [g] ^[65]^.

**Figure 4c** shows the phase trends at 1 kHz under the three temperature conditions. In all cases, the phase change was not significant at the failure point in the 87 °C group, as well as in the stable groups at 37 and 67 °C. The detailed normalized impedance trends at 87 °C are presented in **Figure S15**. The impedance trends of failed devices typically showed stable behavior followed by a steep increase, indicating device failure corresponding to the failure modes categorized in **Figure 4a**. The most common failure mode in the 87 °C group was identified as opening site damage with mechanical tearing, as shown in the microscopy images in **Figure S15**. Other failure modes were further categorized once the channels reached the failure threshold based on impedance change (**Figure S15**).

Capacitance evolution also provides insight into device lifetime and functionality (**Figure 4d**). Similar to impedance trends over time, capacitance may either decrease or increase as an indicator of device failure. A decrease in capacitance is often associated with PEDOT:PSS degradation, such as opening site damage or wiring failure. Conversely, an increase in capacitance may result from encapsulation delamination, which increases the effective surface area beyond the designed opening site. As shown in **Figure 4d**, capacitance gradually decreased toward device failure in the 87 °C group, while devices in the 37 °C and 67 °C groups maintained relatively stable normalized capacitance over time, indicating minimal degradation. In general, capacitance decreased as impedance increased over time (**Figure S16**). This inverse relationship is consistent with loss of effective electrochemically active area due to structural or interfacial degradation.

Interestingly, one major difference between the 67 °C and 87 °C device groups is the surface color transition of the devices from transparent to yellow with visual inspection, as shown in **Figure S17**. Previous studies have reported that exposing fluorinated polymers to high temperatures induces dehydrofluorination, resulting in a color change to yellow or brownish, which was also observed in this study.^[57]^ Devices immersed at 87 °C for 30 days (equivalent to ∼2.63 years at 37 °C) exhibited thermal discoloration that was not observed in devices soaked at 67 °C for 250 days (equivalent to ∼5.47 years) (**Figure S17**). Further XPS results support that the color transition is due to dehydrofluorination, with the transition of C–F groups into carbon-based groups (**Figure S18**). Notably, despite the shorter equivalent aging time at 87 °C, more severe changes were observed compared to the 67 °C group. This indicates that degradation at 87 °C is governed by thermally activated chemical reactions that are not representative of physiological degradation pathways. Furthermore, studies on PEDOT:PSS conductivity have shown that temperature alone significantly affects its electrical properties, with temperatures above 75 °C causing a drastic decrease in conductivity.^[58]^Therefore, accelerated aging at elevated temperatures may introduce degradation mechanisms that differ from those under physiological conditions, limiting the validity of lifetime projection based solely on Arrhenius scaling for organic materials. ^[55]^

**Figure 4e** shows the representative Bode plots of the device before and after stored under 37 and 67 °C for 325 days. As shown here, there was no significant difference between before and after soaking under 1× PBS for 325 days across the overall frequency from 1 Hz to 100 kHz implying stable device performance. **Figure 4f** shows the CV curve trend before and after soaking under 37 and 67 °C 1× PBS. For the electrode stored under 37 °C, there was no significant change in terms of shape and area under the curve implying the stability of the device showing C/C_0_ value of 0.96 and 0.98 for anodic and cathodic value, respectively. For the electrode stored under 67 °C, there was no significant reduction in normalized capacity (0.86 and 0.94 for anodic and cathodic responses, respectively), although slight peak broadening was observed. This behavior may be attributed to subtle structural densification or interfacial changes at the opening site under elevated temperature, which can influence charge transport kinetics without significantly degrading electrochemical capacity. **Figure 4g** shows the average impedance change at 10 Hz, 1 kHz, and 100 kHz at 37 and 67 °C. As shown, the impedance values remained stable within the 10 % variation before and after soaking under 1× PBS for 325 days.

Further analysis included XPS analysis and cross-sectional SEM imaging with EDS characterization of devices soaked under the three temperature conditions were utilized to assess the efficacy of the fluorinated polymer as an ionic barrier. First, XPS analysis was performed to evaluate ion penetration, specifically Na, K and Cl ions enriched within the PBS domain (**Figure 4h** and **Figure S19**-**20**). The results, obtained with argon gas etching from top to the surface to beneath the encapsulation layer, show that ionic peaks were present only at the surface directly exposed to the PBS solution. In contrast, the absence of such peaks throughout the etching depth indicates no significant ion penetration beneath the encapsulation layer, validating its effectiveness as an ionic barrier. Next, SEM-EDS analysis was adopted to evaluate structural stability through morphological analysis and assess ion penetration through the encapsulation or substrate layers (**Figure S21**-**S25**). As shown in **Figure S20**, no significant delamination was observed among the encapsulation, conductive traces, or substrate layers, indicating stable structural integrity. Additionally, no substantial ion penetration (Na, Cl, K) was detected across all temperature groups, with elemental distributions comparable to the pristine device (**Figure S22-S25**). These results confirm the effectiveness of PVDF-HFP as a robust ionic barrier under both physiological and moderately accelerated conditions.

**Figure 4i** shows the comparison of projected lifetimes of previously reported implantable microelectrodes with various materials. The device introduced in this study demonstrates excellent projected longevity compared to previously reported implantable microelectrodes, highlighting its potential as a durable platform for chronic implantable biointerfaces.

### 2.5. In vitro biocompatibility assessment and comparison among materials

For implantable devices, maintaining cell integrity and compatibility with biological systems is essential. In our previous study, the printed PVDF-HFP/PEDOT:PSS sensor fabricated via EHD printing demonstrated excellent cell viability and growth.^[17]^ Additionally, scratch assays of cell cultures on the printed sample platform further validated its safety by showing recovery of cell growth in damaged regions.^[17]^ To further evaluate the biocompatibility of the printed materials, particularly PVDF-HFP, in vitro testing was conducted with direct comparison to widely used biocompatible substrates, polydimethylsiloxane (PDMS) and polyimide (PI). In this assessment, human normal dermal fibroblasts (HNDFs) were cultured on PDMS, PI, and PVDF-HFP substrates for 5 days (**Figure 5a**). Cell morphology and growth patterns were monitored using brightfield imaging on Day 1 and Day 5 (**Figure 5b**). Across all substrates, cells exhibited characteristic fibroblast morphology with clear cellular boundaries, and cell density markedly increased by Day 5, indicating that none of these substrates induced observable cytomorphological abnormalities.

**Figure 5.**
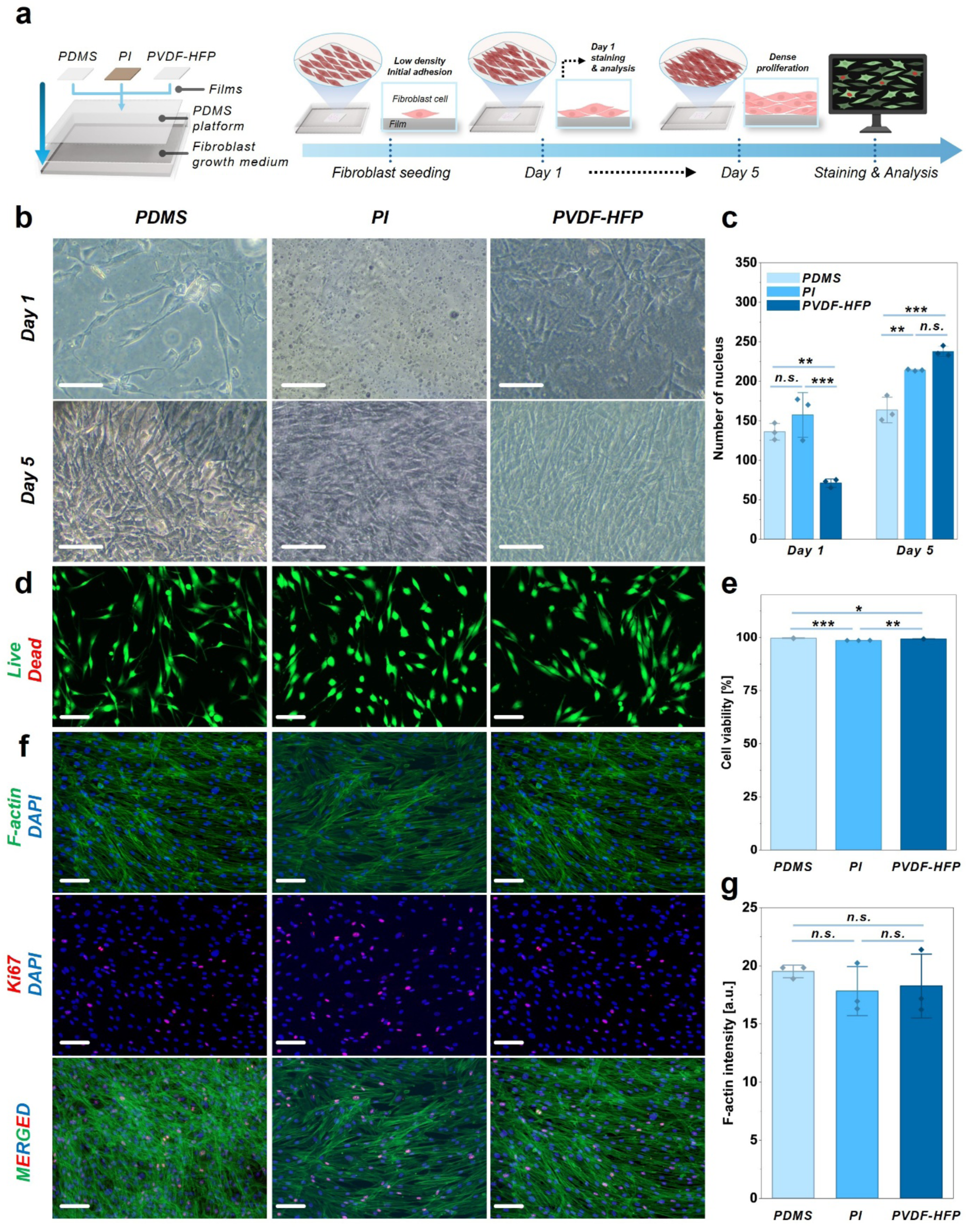
In vitro biocompatibility testing of three different substrates with fibroblast cells. (a) Schematic illustration of experimental design and protocol. (b) Representative bright-field images showing cell morphology and growth patterns across different substrate material groups. (c) Quantitative analysis of DAPI-positive cell counts across groups measured on Day 1 and Day 5. (d) Immunofluorescence staining of Calcein-AM (live, green) and Propidium Iodide (dead, red). (e) Quantitative analysis of the proportion of live cells relative to the total cell population across groups. (f) Immunofluorescence staining of cytoskeletal and proliferative markers; F-actin (green) indicates cytoskeletal organization and cell spreading, and Ki67 (red) indicates proliferative activity, with nuclei counterstained with DAPI (blue). (g) Quantitative analysis of F-actin intensity, indicating differences in cytoskeletal organization across groups. (n = 3). Error bars represent standard deviation. *, p ≤ 0.05; **, p ≤ 0.01; ***, p ≤ 0.001; n.s., not significant. Scale bars: 100 µm.

To quantitatively assess cell proliferation, DAPI-stained nuclei were counted at both time points (**Figure 5c, Figure S26**). On Day 1, the PVDF-HFP substrate supported significantly greater cell attachment compared to PDMS, while no significant difference was observed between PDMS and PI. PVDF-HFP also exhibited significantly higher nucleus counts than PI on Day 1, suggesting that the PVDF-HFP surface facilitates superior initial cell adhesion. By Day 5, both the PVDF-HFP and PI substrates showed significantly greater nucleus counts than PDMS while no significant difference was detected between PVDF-HFP and PI. Overall, the PVDF-HFP substrate achieved the highest cell counts across the entire culture period, indicating that its surface properties are particularly conducive to HNDF adhesion and sustained proliferation.

Cell viability was subsequently evaluated using a Live/Dead assay with Calcein-AM (live, green) and Propidium Iodide (dead, red) staining at Day 5 (**Figure 5d, e**). Although statistically significant differences were detected among groups, cell viability remained consistently high up to 90% across all substrates, and no group exhibited evidence of cytotoxicity. These findings confirm that the PVDF-HFP substrate does not compromise cell survival, supporting its biological safety for use in contact with tissue.

To further characterize the cellular response at the cytoskeletal level, immunofluorescence staining for F-actin and Ki67 was performed at Day 5 (**Figure 5f**). F-actin staining was used to evaluate cytoskeletal organization and cell spreading, as the integrity and arrangement of actin stress fibers reflect the degree to which cells have adapted to and engaged with the underlying substrate. Cells on all three substrates displayed well-organized F-actin filaments with elongated morphology, consistent with healthy fibroblast behavior. Quantification of F-actin fluorescence intensity revealed that cells cultured on PVDF-HFP exhibited significantly higher F-actin intensity compared to PDMS, while F-actin intensity on PVDF-HFP was comparable to that on PI (**Figure 5g**). The comparable F-actin intensity to PI, a widely used biocompatible polymer substrate, indicates that the PVDF-HFP surface supports cytoskeletal organization and cell spreading at a similar level. In parallel, Ki67 staining was performed as a marker of active cell proliferation. Ki67-positive nuclei were consistently observed across all substrate groups throughout the 5-day culture period, confirming that none of the substrate materials suppressed cell cycle progression or induced growth arrest, further supporting the biocompatibility of the PVDF-HFP substrate.

### 2.6 In vivo biocompatibility assessments for printed soft microelectrode

In addition to the in vitro biocompatibility test, in vivo biocompatibility study was conducted with the subcutaneous implantation model in BALB/c rodents. The aim of this study was to evaluate the bio-integration characteristic of PVDF-HFP/PEDOT:PSS-based printed soft microelectrodes in terms of collagen formation, inflammatory response, and fibrotic signaling by comparing widely used biocompatible materials under implanted condition.

**Figure 6a** shows the schematic illustration of the subcutaneous implantation procedure. For this study, three types of miniaturized printed patches were prepared: PDMS/PEDOT:PSS, PI/PEDOT:PSS, and PVDF-HFP/PEDOT:PSS. These samples were implanted subcutaneously for two weeks, after which tissue responses were analyzed with sham group (**Figure 6b**). After two weeks of implantation, fibrous capsule formation was observed at the implantation sites in all groups, as shown in hematoxylin and eosin (H&E) staining results (**Figure 6c**). Further evaluation using Masson’s trichrome (MT) staining was performed to assess collagen deposition, which is a key factor in capsule formation. As shown in **Figure 6c**, the PDMS group exhibited the largest collagen formation (bright blue) area among the patch groups, while the PI and PVDF-HFP groups showed relatively smaller collagen deposition compared to the PDMS and sham groups, with a trend toward reduced fibrotic response. Inflammatory response was evaluated using CD68 immunostaining (**Figure 6c**). The results indicate that the PVDF-HFP group did not show significant macrophage infiltration compared to the other groups.

**Figure 6.**
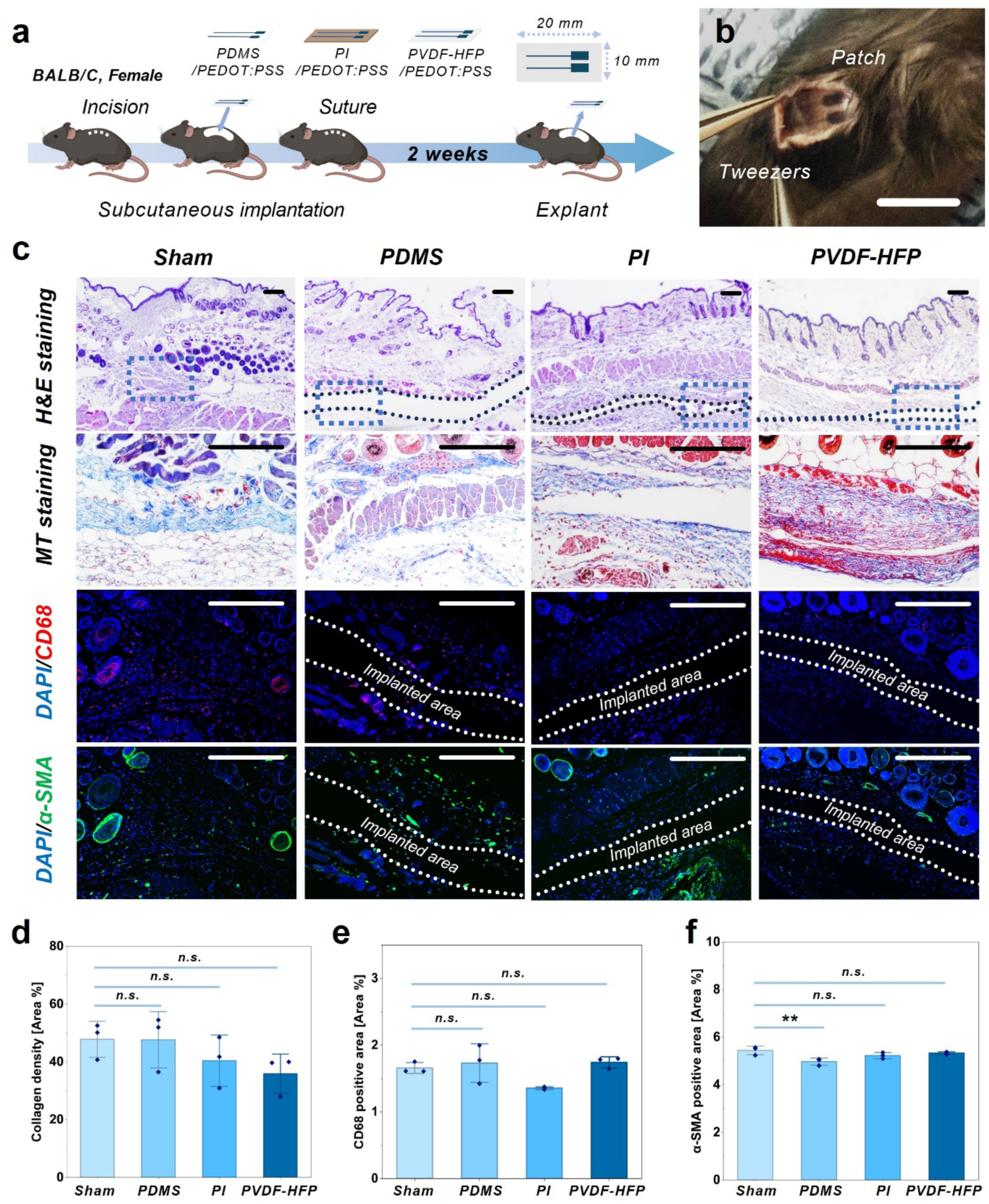
In vivo biocompatibility testing with subcutaneous implantation. a) Schematic illustration of the experimental design and surgical protocol. b) Representative photograph of the subcutaneously implanted printed device (scale bar: 1 cm). c) Representative histological and immunofluorescence analyses of regenerated tissues, where hematoxylin and eosin (H&E) staining shows overall tissue morphology and epithelial restoration, and Masson’s trichrome (MT) staining indicates collagen deposition (blue), reflecting fibrotic changes. Immunofluorescence staining demonstrates CD68-positive macrophages (red) and α-SMA positive myofibroblasts (green), with nuclei counterstained with DAPI (blue), and dashed boxes indicate regions of interest (scale bars: 200 µm). d) Quantification of collagen deposition based on MT staining, indicating the extent of fibrosis within regenerated tissues. e) Quantitative analysis of histological and immunofluorescence data, including inflammation, fibrosis, and tissue regeneration across groups. f) Immunofluorescence analysis of inflammatory and fibrotic markers, where sections were stained for CD68 (red) and α-SMA (green), with DAPI (blue) marking nuclei, indicating macrophage infiltration and myofibroblast activation, respectively. Error bars representing the standard deviation (n = 3). **, p ≤ 0.01; n.s., not significant.

Further, immunofluorescence staining for α-SMA, a marker of myofibroblasts, was performed to assess fibrotic activity around the implantation site (**Figure 6c**). Fibrous capsule formation is generally associated with myofibroblast activation, which is often attributed to mechanical mismatch between the tissue and implanted material.^[66]^ As shown in **Figure 6c**, no significant α-SMA expression was observed in the PVDF-HFP group compared to the PDMS group, which has a lower elastic modulus and is likely to have better integration with biological system. This suggests that PVDF-HFP provides comparable tissue compatibility to well-established soft materials. **Figure 6d** presents the quantitative analysis of collagen formation across all groups. The results indicate no significant differences among the groups under statistical analysis. Notably, the PVDF-HFP group showed the lowest average value, suggesting a tendency toward reduced capsule formation (**Figure 6d**). Similarly, quantitative analysis of CD68 staining showed no significant differences among the groups, indicating comparable inflammatory responses and overall tissue compatibility (**Figure 6e**). The quantification of α-SMA indicated that PDMS had the lowest average value, likely due to its lower stiffness and better mechanical matching with tissue. In contrast, the PVDF-HFP group showed comparable levels of myofibroblast activation, supporting its favorable biocompatibility compared to well-known biocompatible materials (**Figure 6f**).

## 3. Conclusion

The combination of an advanced additive manufacturing method, EHD printing, and functional materials enabled the fabrication of a long-lasting soft microelectrode to investigate mechanical stability, long-term reliability, and biocompatibility. This advanced printing technology allows scalable device fabrication with an integrated multilayer structure, ensuring reliable device performance. The transparent conducting polymer PEDOT:PSS was used as the conductive material, while the fluorinated polymer PVDF-HFP was used as the encapsulation material. EHD-printed PEDOT:PSS demonstrated excellent electrochemical stability under biomimicking conditions. Additionally, the EHD-printed fluorinated soft polymer PVDF-HFP exhibited outstanding performance as a soft encapsulation layer, showing no cracking or delamination under continuous mechanical deformation. Accelerated lifetime testing confirmed the excellent stability of the printed microelectrode. In this test, we found that accelerated aging at temperatures above a certain threshold for soft organic materials may not be suitable for accurately estimating or predicting failure time and mechanisms, as it can induce thermally activated degradation that is not representative of physiological conditions. Furthermore, XPS and SEM-EDS results demonstrated the effectiveness of EHD-printed PVDF-HFP as an ionic barrier, with no significant ion penetration over time. Biocompatibility assessment using fibroblast cells showed excellent cell growth and no signs of cytotoxicity compared with widely used substrate materials, supporting its suitability as an implantable material. In vivo biocompatibility evaluation using a rodent subcutaneous model with different substrate materials and PEDOT:PSS further confirmed its comparable performance in an implanted environment without adverse tissue response. With further design optimization and scaling strategies, this combination of printing technology and materials has strong potential as a next-generation platform for durable, high-density, and miniaturized soft implantable biointerfaces.

## Experimental Section/Methods

### Ink fabrication and batch printing process for five channel microelectrodes

The fabrication of five-channel microelectrodes began with the preparation of a Poly(vinylidene fluoride-co-hexafluoropropylene)/dimethylformamide (PVDF-HFP/DMF) solution (Sigma-Aldrich, USA), where PVDF-HFP pellets (6 g) were dissolved in DMF (40 mL) using a magnetic stirrer for 1 h. Commercially available PEDOT:PSS ink (PK-246T, J. Chem, South Korea) was used for the conductive layer. A schematic illustration of the printing process is shown in **Figure S1**. Prior to printing, the heat bed, which is integrated into an x–y translational stage, temperature was set to 40 °C for preconditioning. A polyethylene terephthalate (PET) film, used as a sacrificial substrate, was placed on the preconditioned printing bed and secured via vacuum assistance. The PVDF-HFP substrate layer was printed three times, followed by three layer of PEDOT:PSS printing to form the conductive traces, and subsequently PVDF-HFP encapsulation layer was printed on top of the PEDOT:PSS traces. The nozzle diameters for PVDF-HFP and PEDOT:PSS were 200 μm and 70 μm (ceramic nozzle, Cosma, South Korea), respectively. For PVDF-HFP (substrate and encapsulation) printing, the printing speed, stand-off height, and nozzle voltage were set to 10 mm s^−1^, 20 μm, and 2 kV, respectively, while for PEDOT:PSS printing these parameters were 20 mm s^−1^, 20 μm, and 0.5 kV. After printing, the heat bed temperature was increased to 80 °C for solvent evaporation and crosslinking of PVDF-HFP. A total of 21 electrodes were printed on a single PET film. After the devices were detached, devices were wired out silver wires (AGT 0525, World Precision Instruments) though connector parts using silver paste and encapsulated with silicone. The total net printing time for one electrode batch (21 electrodes) was approximately 3.5 hours.

### Transmittance measurement for films

Five PVDF-HFP films with varying numbers of printed layers, four PVDF-HFP/PEDOT:PSS films with different numbers of PEDOT:PSS printing passes on a PVDF-HFP trilayer substrate, and a PET film as a control sample were evaluated in this study. The optical transmittance of the samples was measured under ambient (air) conditions using a spectrophotometer (Cary 5000, Varian, USA). The samples were exposed to light in the wavelength range of 400–800 nm, with a 1 mm aperture applied using the sample holder.

### Electrochemical characterization of devices

EIS, CV, and VT measurements were performed using a three-electrode configuration, with Ag/AgCl, platinum, and the device serving as the reference, counter, and working electrodes, respectively. Data acquisition was carried out using a CHI 660E electrochemical workstation (CH Instruments, Bee Cave, TX, USA). During testing, the device was fully immersed in 1× phosphate-buffered saline (PBS), and the temperature was maintained at 37 °C using a heated water bath (Corning Inc., Corning, NY, USA). For EIS measurements, a voltage amplitude of 0.05 V was applied. The charge values from the CV curves were calculated by integrating the current over time using MATLAB. The charge storage capacity (CSC) and charge injection capacity (CIC) were calculated using the following equations:

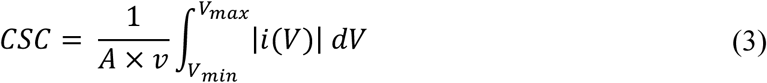

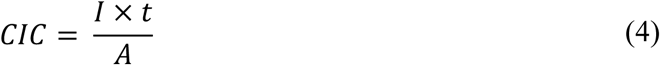

, where *A*, *v*, *Vₘₐₓ*, and *Vₘᵢₙ* denote the electrode area, scan rate, and voltage limits, respectively; *i(V)* represents the current as a function of voltage; and *I* and *t* denote the applied current and time, respectively.

### Measurement of device mechanical properties

For the mechanical characterization of the printed samples, a micro-tensile tester (Modular Force Stage, Linkam Scientific Instruments, UK) with a 200 N load cell was utilized. During tensile testing, both ends of the samples were fixed with clamps and uniaxially stretched at a rate of 100 μm s^−1^, while data were recorded using the OEM LINK software (Linkam Scientific Instruments, UK). Images for observing crack propagation in the devices were captured using an optical microscope (Eclipse LV100N POL, Nikon, Japan) with a FLIR Grasshopper 3 camera (GS3-U3-32S4C-C).

### Monotonic stretching of device under PBS with manual stage

Both ends of the sample were fixed to a manual stage and uniaxially stretched from 0 to 10% strain with 2% increments while immersed PBS, and the corresponding impedance values were measured. The temperature of the PBS was maintained at 37 °C, consistent with the conditions used in the initial electrochemical characterization of the device.

### Cyclic tensile testing under PBS

Cyclic tensile testing was performed under phosphate-buffered saline (PBS) using a micro-tensile tester (Modular Force Stage, Linkam Scientific Instruments, UK) with a 200 N load cell. Prior to testing, PBS was filled into a temperature-controlled reservoir with a microheater and preheated to 37 °C. The device was then fixed at both ends using clamps and subjected to 1,000 cycles of 1% strain. The impedance of the device was measured before and after cyclic loading. For image acquisition, an optical microscope and camera were used, following the same setup as described for mechanical testing.

### Vial preparation for lifetime testing and lifetime assessments

Vial preparation began by drilling five holes into the vial cap to allow the electrode wires to pass through for data collection (**Figure S13**). The wires extending from the cap were secured using adhesive epoxy (Loctite, USA). The vial was then filled with 1× PBS up to a level below the connector to simulate implanted conditions. Due to hydrophobicity of the device, the metal clip was mounted at the end of the sample to be fully soaked under the PBS. The prepared devices were stored in ovens set at three different temperatures for accelerated aging. EIS and CV measurements to assess the device performance during the lifetime testing were conducted under the same conditions as those used for the initial electrochemical characterization of the devices.

### XPS analysis for ion penetration assessments and dehydrofluorination of PVDF-HFP film

X-ray photoelectron spectroscopy (XPS) was performed to analyze the surface chemical composition and evaluate the penetration of Na, Cl, and K species after exposure to 1× PBS under different temperatures and soaking durations. A Thermo Fisher Scientific spectrometer (K-Alpha, 1200505) equipped with a monochromatic Al Kα X-ray source (1486.6 eV) was used. Sample charging was mitigated using an electron flood gun. The X-ray spot size was set to 100 μm, the take-off angle was 90°, and the analyzer vacuum was maintained at least 5.0 × 10^−8^ mbar prior to sample processing. Depth profiling was carried out using argon ion etching with an energy of 3 keV and a sputter rate of 10.62 nm s^−1^ based on a Ta_2_O_5_ standard, to evaluate the subsurface distribution of ions. The etching depth was controlled as a function of time, with 10 incremental steps of 5 s etch time per step from the surface. Spectra were collected after each etching step to monitor variations in Na, Cl, and K signal intensities as a function of depth from the top PVDF-HFP encapsulation layer. For the surface chemical bonding group analysis of PVDF-HFP films before and after soaking at 87 °C, the same scanning parameters were used without an etching process.

### Cell Culture

Human normal dermal fibroblasts (HNDFs) were cultured in Fibroblast Growth Medium (FGM) under standard culture conditions at 37°C in a humidified atmosphere containing 5% CO₂. The culture medium was replaced every 2–3 days, and the cells were subcultured upon reaching appropriate confluency.

### In vitro cell viability assay

Prior to cell seeding, a cell culture platform was prepared by pouring 2 g of PDMS prepolymer mixed with curing agent (10:1, w/w) into a 100 mm dish, followed by curing in a dry oven at 80°C for 1 h. After curing, the PDMS substrate was cut into 1 cm × 4 cm pieces, and the devices were placed on the surface. Cell viability was quantified using a live/dead viability/cytotoxicity assay kit (Sigma-Aldrich, USA). After removing all culture media, 1 µM ethidium homodimer-1 (dead cell indicator; Invitrogen, Sigma-Aldrich, USA) and 5 µM calcein-AM (live cell indicator; Sigma-Aldrich, USA) were prepared in 2 mL of Dulbecco’s phosphate-buffered saline (DPBS) following the method described in our previous study.^[17]^ The cell culture platforms were incubated with the staining solution at 37°C in a 5% CO₂ incubator for 30 min, followed by three washes with DPBS. Fluorescence images were acquired using a confocal laser scanning microscope (QUADSCAN, Live Cell Instrument, Republic of Korea), and cell viability was calculated as the ratio of live cells to the total number of cells. The prepared platforms were sterilized under UV light for 5 min, followed by coating with 0.01% poly-L-lysine solution (Sigma-Aldrich, USA) at 37°C for 10 min. The platforms were then rinsed three times with phosphate-buffered saline (PBS). Subsequently, human normal dermal fibroblasts (HNDFs) were seeded at a concentration of 0.1 mg/mL onto both platforms with and without the collagen I (100 µg/mL; 5225, Advanced BioMatrix) sensor. The cells were cultured for 5 days in a humidified incubator at 37°C with 5% CO₂.

### Fabrication process for the miniaturized patch for in vivo biocompatibility testing

For subcutaneous implantation in a rodent model, three types of samples were prepared: PDMS/PEDOT:PSS, PI/PEDOT:PSS, and PVDF-HFP/PEDOT:PSS. The final dimensions of each patch were 20 mm × 10 mm. The fabrication and printing procedures followed the same protocol as described for the five-channel microelectrode fabrication. For the PDMS/PEDOT:PSS patch, the substrate was prepared by mixing the base and curing agent (Sylgard 184, Dow Corning Corporation, USA) at a 10:1 ratio. After degassing, 10 cc of PDMS was spin-coated onto a PET film for 10 min and cured at 80 °C for 2 h. The cured PDMS substrate was then treated with oxygen plasma (20 W power, duty ratio of 64, and 30 sccm gas flow) for 1 min. The plasma-treated PDMS substrate was transferred to the printer, and the PEDOT:PSS conductive layer was printed using the same parameters as those used for the PVDF-HFP-based microelectrodes. Subsequently, the opening sites were masked, and an additional PDMS layer was coated and cured at 80 °C for 2 h, resulting in a total device thickness of approximately 450 μm. For the PI/PEDOT:PSS samples, the process began by cleaning the surface of the PI film (SKC, South Korea; 75 μm thickness) with isopropyl alcohol (IPA), followed by drying at 40 °C for 10 min. The PI film was then transferred to the EHD printing stage, and the PEDOT:PSS layer was printed using the same parameters as those used for the PVDF-HFP-based microelectrodes. Finally, a single-sided adhesive PI film (Alphaflon, South Korea; 65 μm thickness) was laser-cut and attached to the interconnect region.

### In vivo subcutaneous the device implantation

Seven-week-old female BALB/c mice were obtained from Orient Bio (Republic of Korea). The animals were housed under standard laboratory conditions with a 12 h light/dark cycle at room temperature (20–22°C) and provided ad libitum access to food and water. For subcutaneous implantation, mice were anesthetized with isoflurane administered via an oxygen inhalation anesthesia system. The dorsal region was selected as the surgical site, and a 1 cm incision was made on the back. The skin layers, including the epidermis, dermis, and stratum corneum, were carefully dissected to expose the subcutaneous space at the designated implantation site. PDMS, PVDF, and PI film materials were implanted into the prepared space, with two samples placed on each side per mouse. The control group underwent the same surgical procedure without implantation of materials. After 14 days, the mice were euthanized, and the implanted tissues were harvested for subsequent in vivo analysis. All animals were maintained under specific pathogen-free conditions at the Laboratory Animal Center of Sungkyunkwan University and handled in accordance with institutional guidelines approved by the Institutional Animal Care and Use Committee (IACUC) of Sungkyunkwan University (SKKUIACUC2025-05-50-1).

### Histology (H&E, Masson trichrome staining)

All tissue samples were fixed in 4% paraformaldehyde (PFA) overnight and subsequently embedded in 2% agarose to stabilize the tissue structure, followed by solidification. The samples were then transferred to 70% ethanol for initial dehydration. Tissues were further dehydrated through a graded ethanol series (80–100%) and subsequently cleared with Histoclear (HS-202, National Diagnostics, Charlotte, NC, USA) for 2 h to remove residual ethanol. After clearing, tissues were thoroughly infiltrated with paraffin, embedded in molds filled with molten paraffin, and allowed to solidify. The paraffin-embedded tissues were then sectioned into 5 μm-thick slices and mounted onto glass slides. Paraffin-embedded tissue sections (5 μm) were deparaffinized using Histoclear (National Diagnostics, USA) and subsequently rehydrated through a graded ethanol series from 100% to 70%. The sections were stained using hematoxylin and eosin (H&E) and Masson’s trichrome staining kits (Agilent, USA) following the method described in our previous study.^[67]^

### Immunostaining

Prior to staining, sections were deparaffinized using Histoclear and rehydrated through a descending ethanol gradient (100% to 80%). Antigen retrieval was performed in sodium citrate buffer (pH 6.0) at 95°C for 30 min using a microwave. Following washing steps, hydrophobic barriers were created around the tissue sections using a DAKO pen to confine the staining area. The samples were then blocked with 5% bovine serum albumin (BSA) in PBS at room temperature for 1 h and subsequently incubated overnight at 4°C with primary antibodies against CD68 (ab53444 abcam, Cambridge, UK, 1:200)and α-SMA(MA1-06110 abcam, Cambridge, UK, 1:200). After washing, the samples were incubated with fluorescent secondary antibodies, including anti-rabbit IgG (Invitrogen, Carlsbad, CA, USA) or anti-rat IgG (Invitrogen, Carlsbad, CA, USA), at a dilution of 1:500 for 1 h at room temperature. Nuclei were then counterstained with DAPI (62248, Thermo Fisher, Waltham, MA, USA; 1:1000) for 10 min. Finally, the samples were mounted using mounting medium (S3025, DAKO, Glostrup, Denmark) and covered with a coverslip for subsequent imaging and analysis following the described method in our previous study.^[68]^

## Supporting Information

Supporting Information is available from the Wiley Online Library or from the author.

## Author Contributions

H.J. and G.L. contributed equally to this work. Conceptualization: H.J., J.-H.S, and K.K. Device design/fabrication: H.J., Y.S., M.K, J.-H.S., and K.K. Methodology: H.J., G.L., Y.S., S.Y.K., and R.M. Investigation: H.J., G.L., Y.S., S.Y.K., R.M., D.C., A.A., S.L.S., and K.P. Visualization: H.J. and G.L. Supervision and funding acquisition: J.A., J.-H.S., and K.K. Writing-original draft: H.J. and G.L. Writing-review and editing: H.J., G.L., J.A., J-H.S., and K.K. All authors have read the manuscript and provided comments.

## Acknowledgements

This work was supported by the National Research Foundation of Korea (RS-2024-00347831, J.-H.S., K.K.; RS-2026-25497630, RS-2024-00350442, J.A.) and the Korea Health Technology R&D Project through KHIDI, funded by the Ministry of Health & Welfare, Republic of Korea (RS-2024-00406054, J.A.). This work was also supported in part by Blackrock Neurotech/Blackrock Microsystems Inc. (K.K.) and by the US Army DEVCOM Ground Vehicle Systems Center (GVSC) under Cooperative Agreement W56HZV-21-2-0001 through the VIPR-GS program (K.K.). DISTRIBUTION STATEMENT A. Approved for public release; distribution is unlimited. OPSEC10640

## Data Availability Statement

The data that support the findings of this study are available from the corresponding author upon reasonable request.

