## Supporting Information for "Long-Lasting Electrohydrodynamically Printed Transparent Soft Microelectrode for Implantable Biointerfaces"

†Equally contributed

*Corresponding author

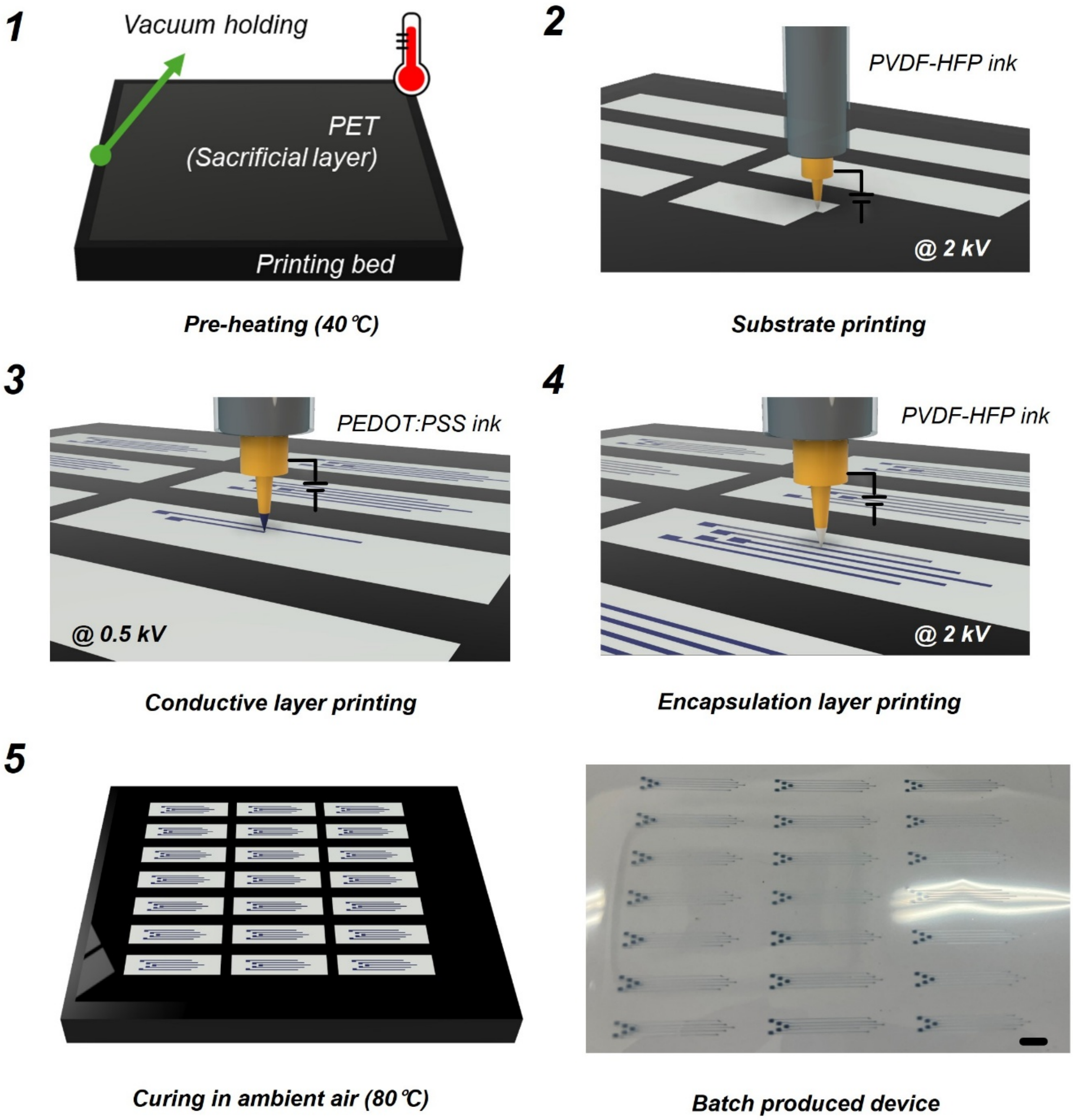


**Figure S1.** Schematic illustration of the fabrication process and photograph image of batch-produced devices. Scale bar: 1 cm.

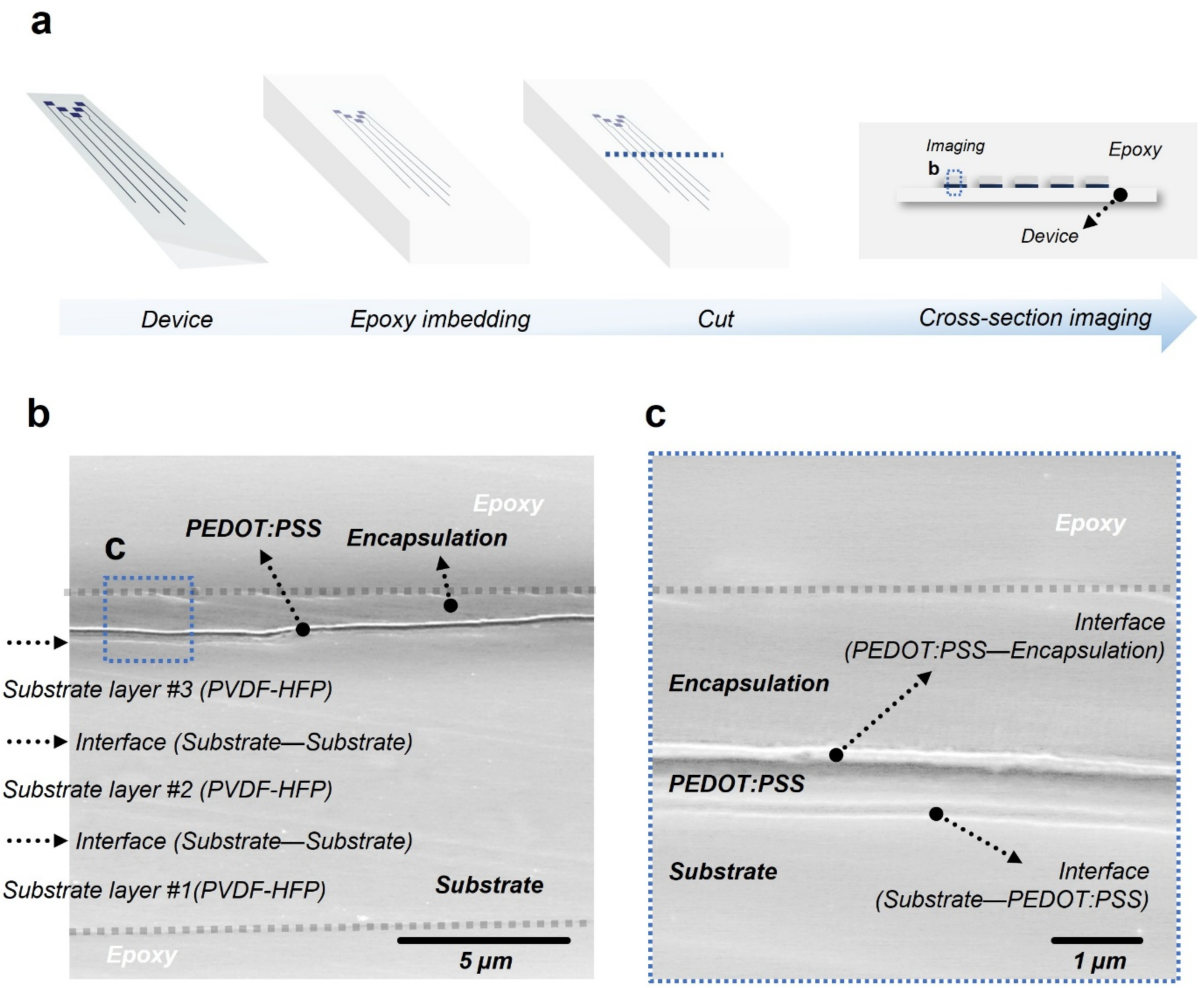


**Figure S2.** a) Schematic of the sample cross-sectioning process. Representative cross-section SEM images of post-fab sample: b) overall view and c) magnified view of the encapsulation–conductive layer–substrate interfaces.

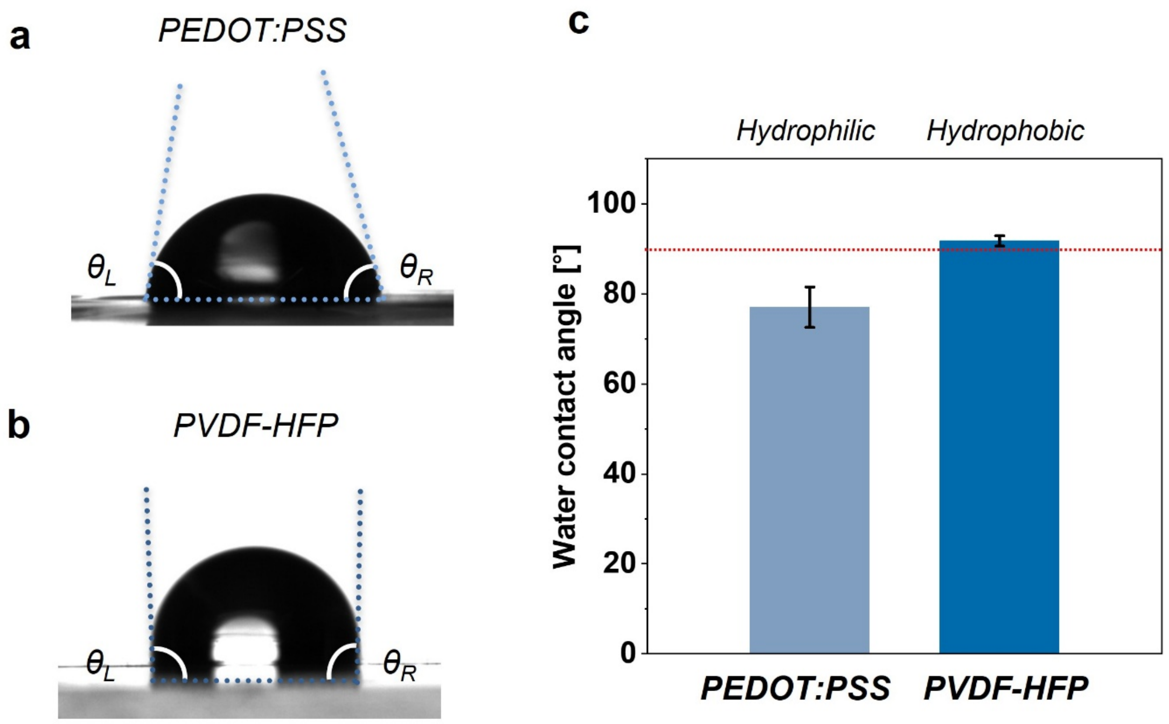


**Figure S3.** Water contact angle testing. Photograph of water droplets on a) the PEDOT:PSS layer and b) the PVDF-HFP layer. c) Water contact angle measurement results (n = 3, averaged from left and right contact angles).

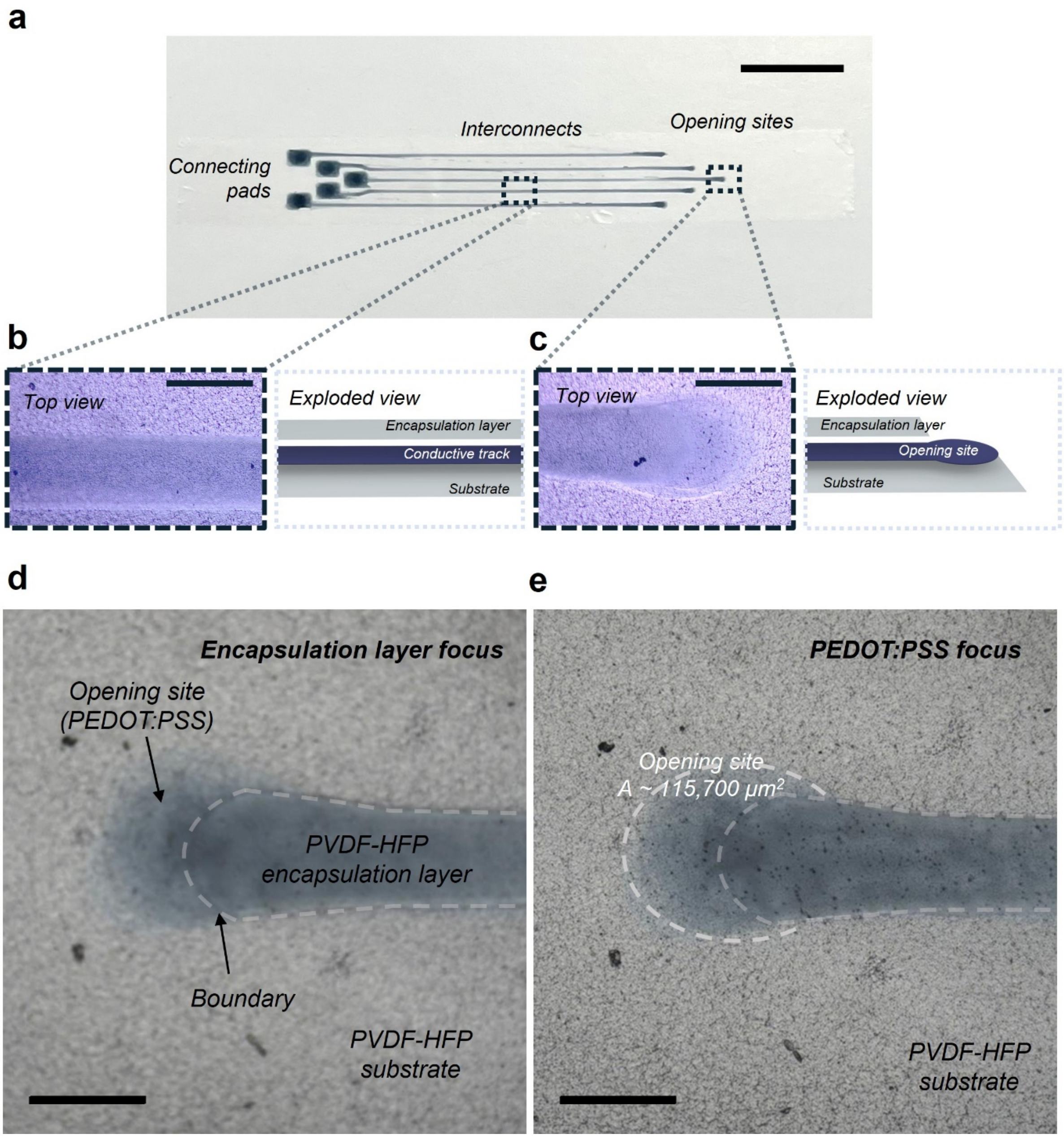


**Figure S4.** a) Photograph of the device. Scale bar: 1 cm. b–c) Optical microscopic images (left) and schematic illustrations (right) of the interconnect (b) and opening site (c). The optical microscopic images shows the images from the top view and schemtaitc illustration show the exploded view of the structure. Scale bars: 300 µm. d–e) Optical microscopic images of the opening site focused on d) the encapsulation layer and e) the PEDOT:PSS layer. Scale bars: 300 µm.

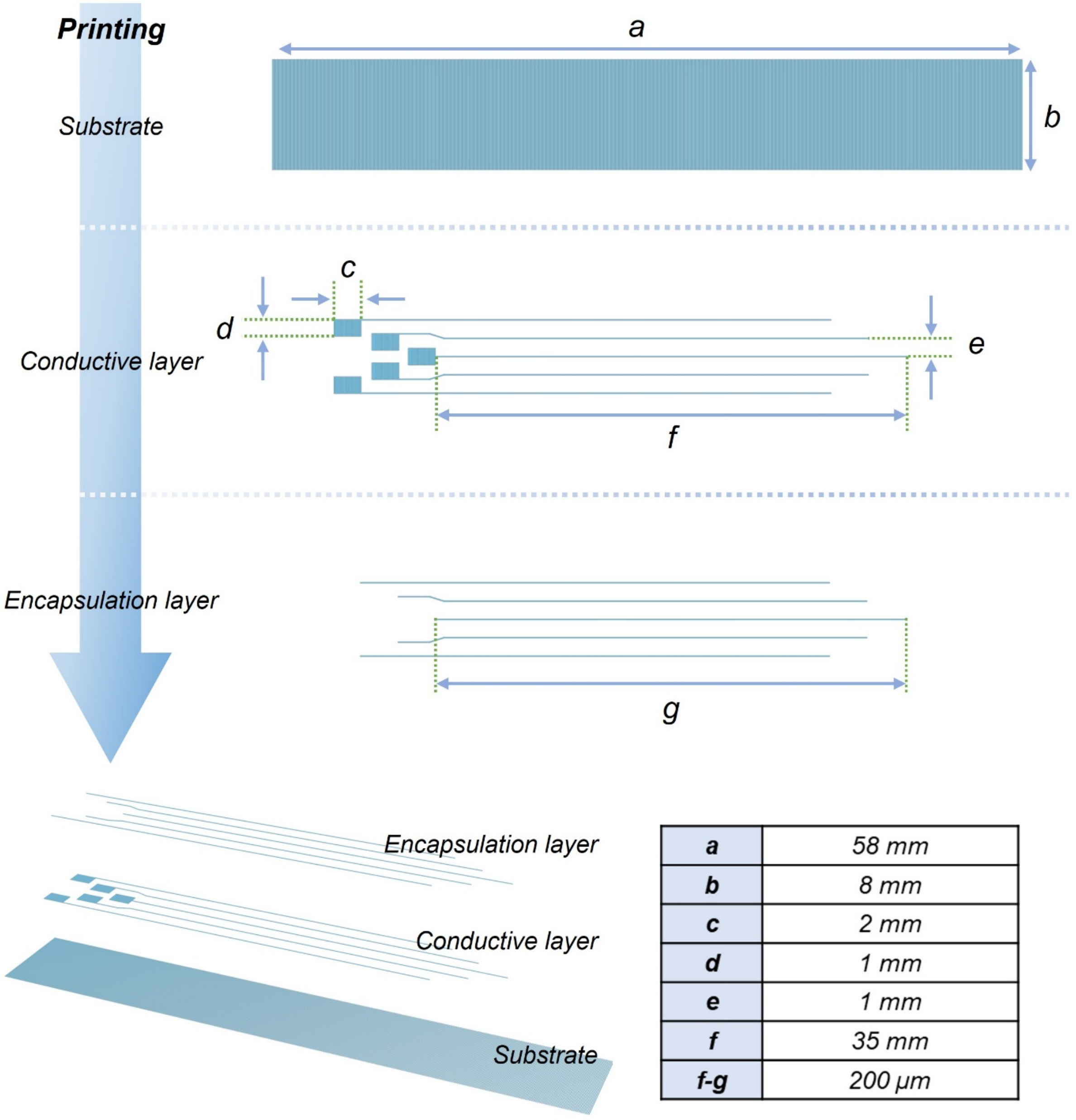


| | |
|---|---|
| **a** | *58 mm* |
| **b** | *8 mm* |
| **c** | *2 mm* |
| **d** | *1 mm* |
| **e** | *1 mm* |
| **f** | *35 mm* |
| **f-g** | *200 µm* |

**Figure S5.** CAD design parameters of the device.

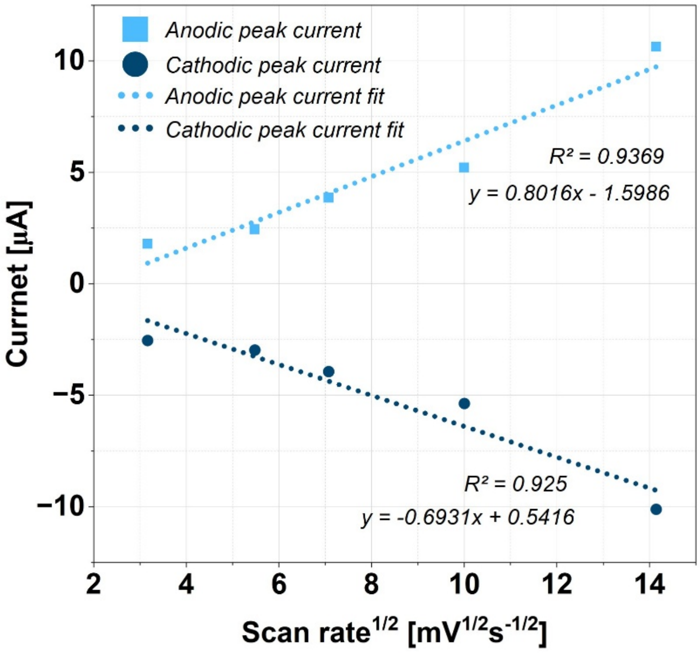


**Figure S6.** Anodic and cathodic peak currents under various scan rates.

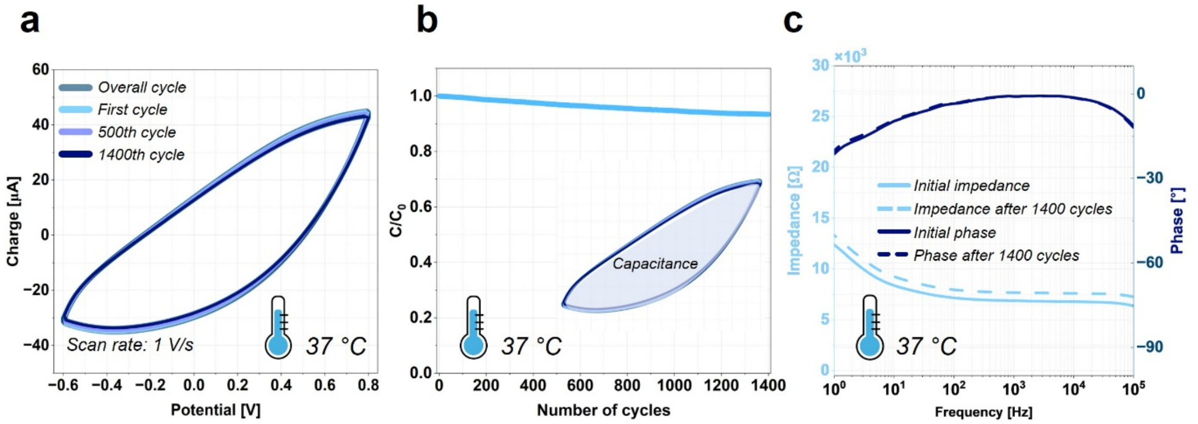


**Figure S7.** CV aging test. a) CV testing results over 1400 cycles. b) Capacitance change during cyclic testing. c) Bode plots of the device before and after CV aging with 1400 cycles**.**

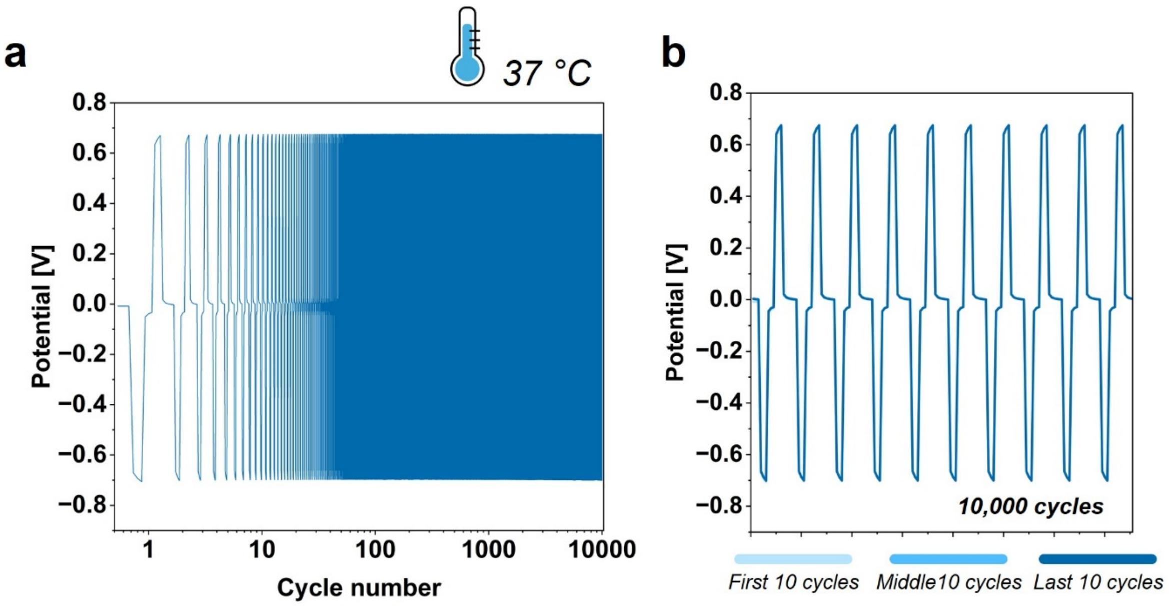


**Figure S8.** Voltage transient cyclic testing. a) Overall 10,000 cycles. b) First, middle, and last 10 cycles.

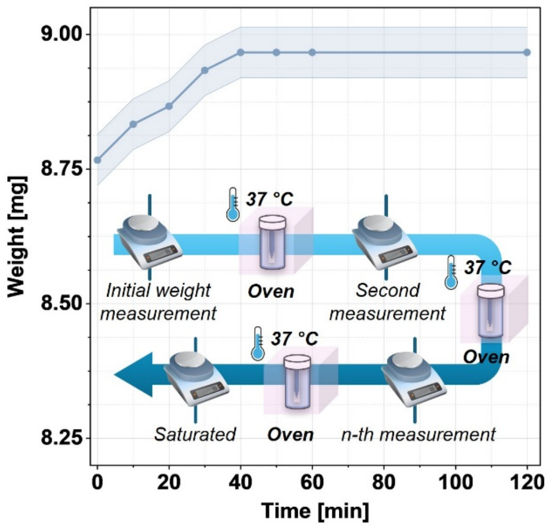


**Figure S9.** Schematic illustration of methods for sorption test and results.

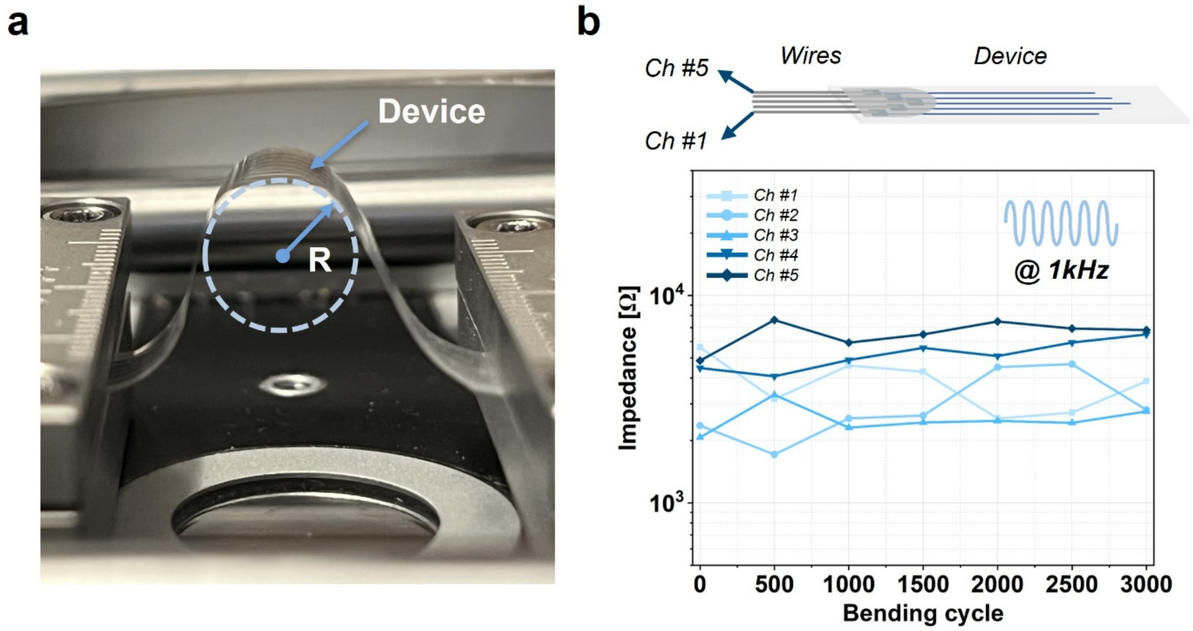


**Figure S10.** Mechanical bending testing of the device. a) Photograph of the testing setup and b) impedance change of five channels over bending cycles. Inset: schematic illustration of the device with wires.

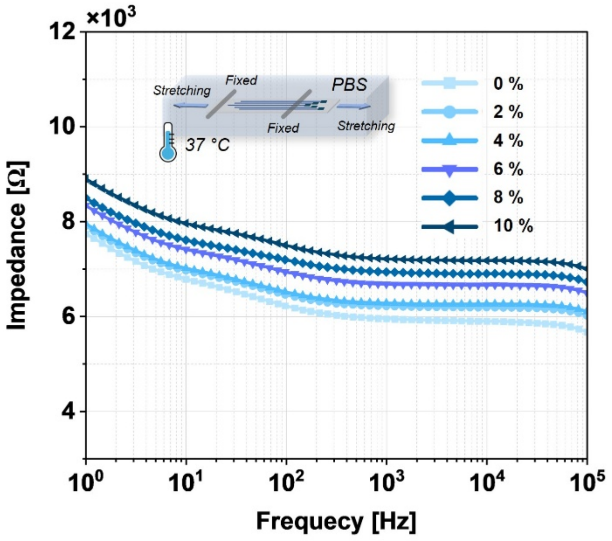


**Figure S11.** Impedance change across a frequency range of $10^0$ to $10^5$ Hz under applied strain in 1× PBS at 37 °C. Inset: schematic illustration of the testing condition.

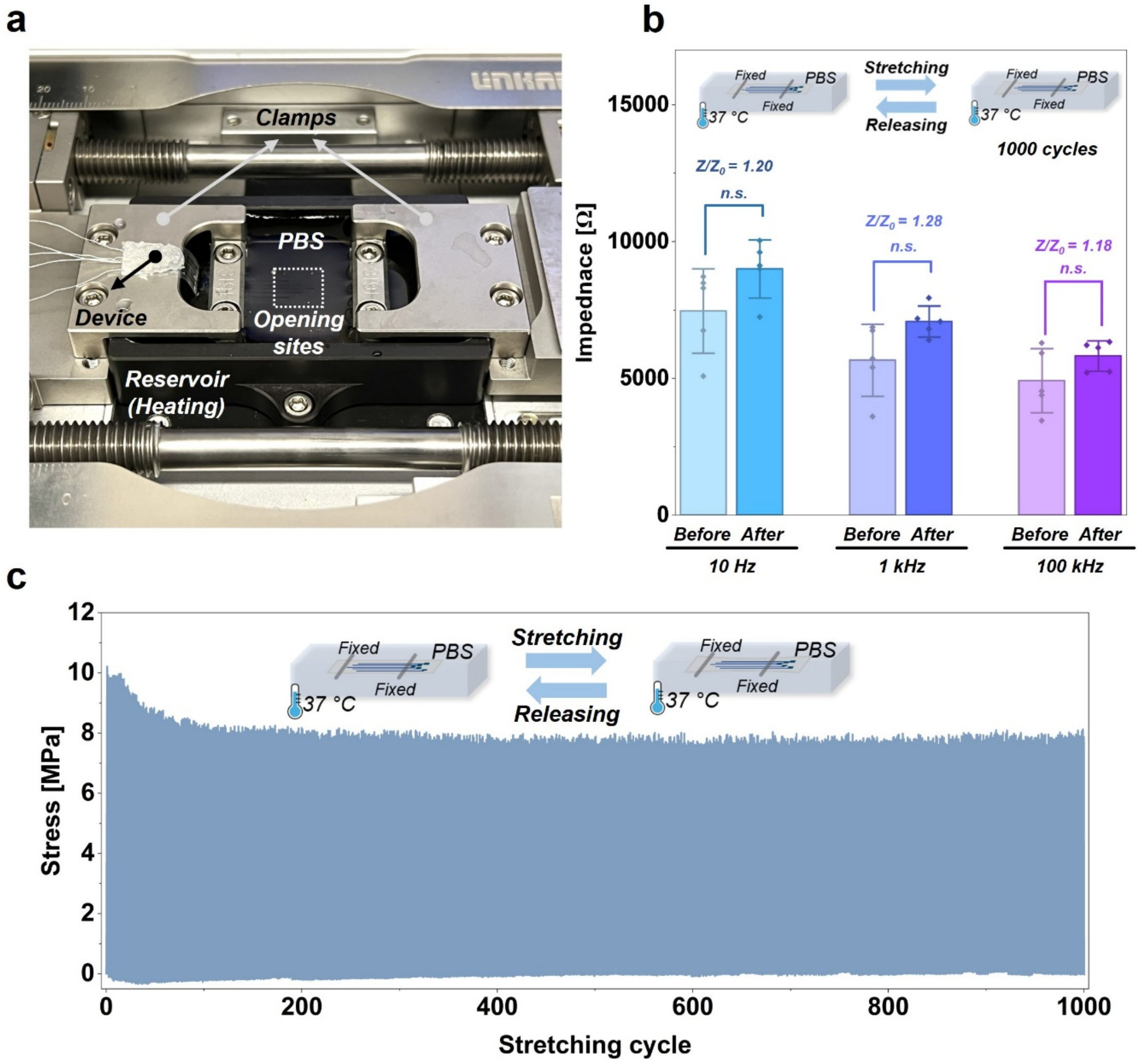


**Figure S12.** Cyclic loading test (1,000 cycles) under 1× PBS at 37 °C. a) Photograph of the testing setup. b) Impedance at 10 Hz, 1 and 100 kHz before and after mechanical cyclic loading in PBS (n=5). c) Stress–strain cyclic plot (applied strain: 1%). Inset: schematic illustration of the testing condition. n.s., not significant.

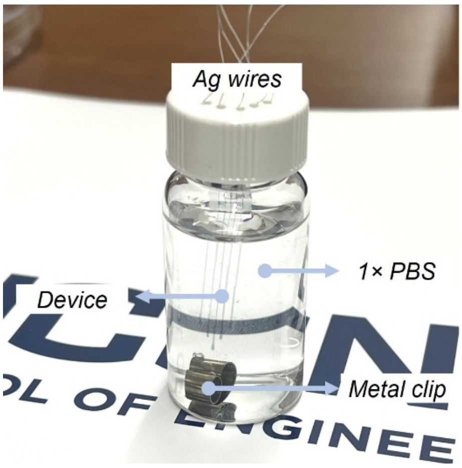


**Figure S13.** Photograph image of sample vial for the lifetime testing.

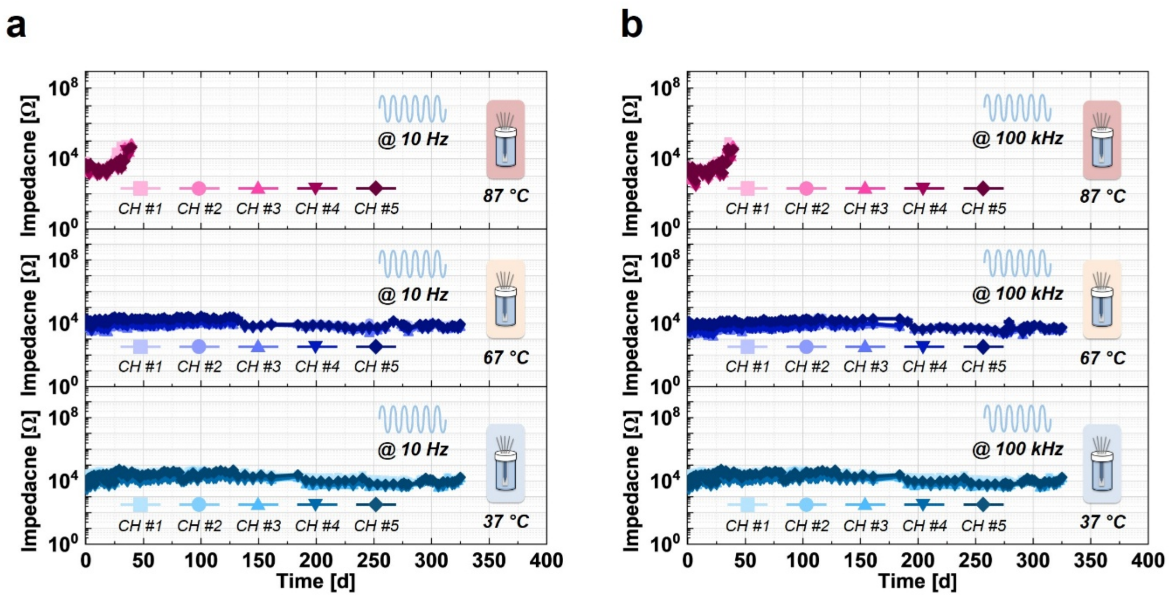


**Figure S14.** Impedance change over time under 1× PBS at different temperatures and frequencies: a) 10 Hz, b) 100 kHz.

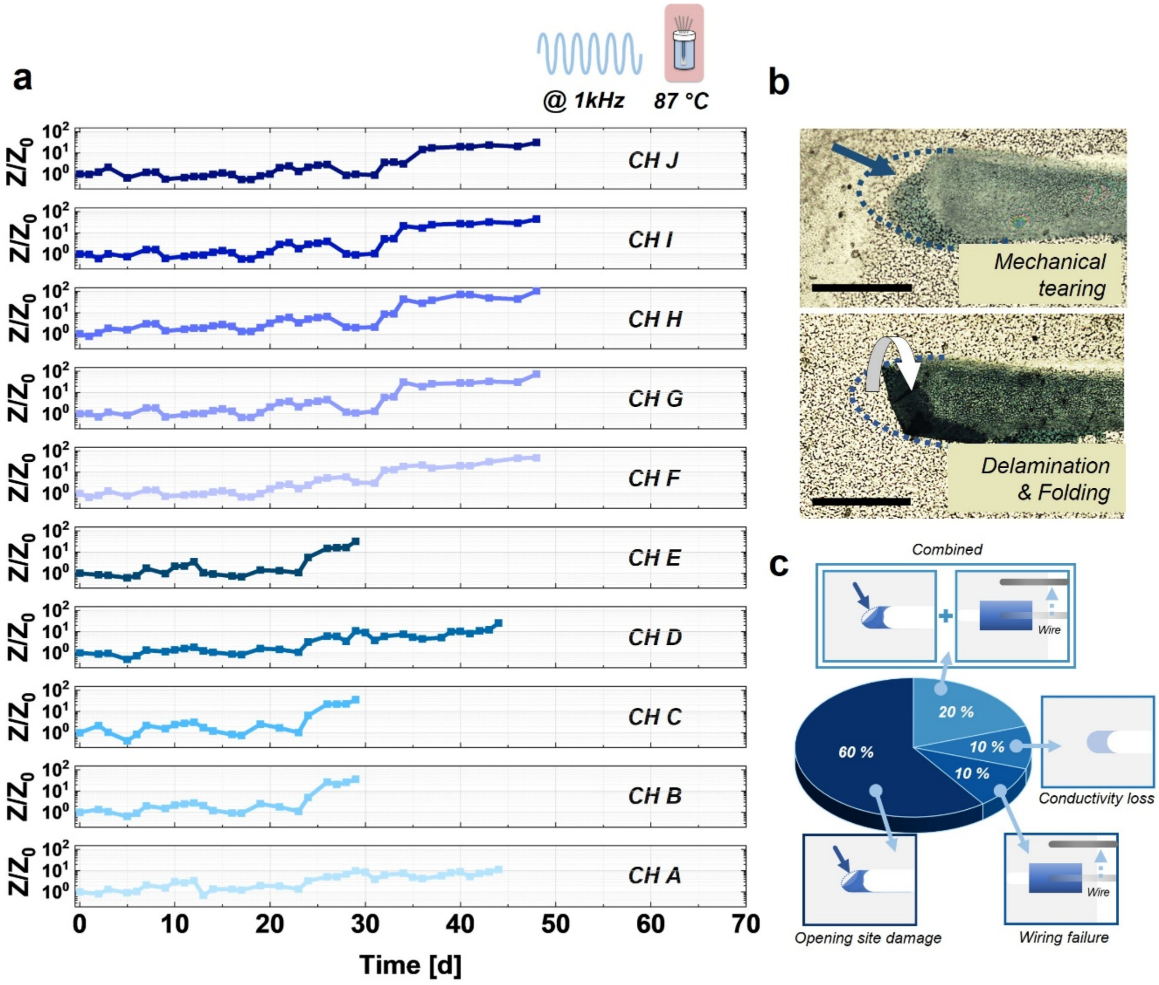


**Figure S15.** a) Normalized impedance change over time at 1 kHz under 87 °C in 1× PBS. b) Optical microscopic images of representative opening-site damage failure modes (scale bars: 200 μm). c) Failure mode distribution of devices tested at 87 °C.

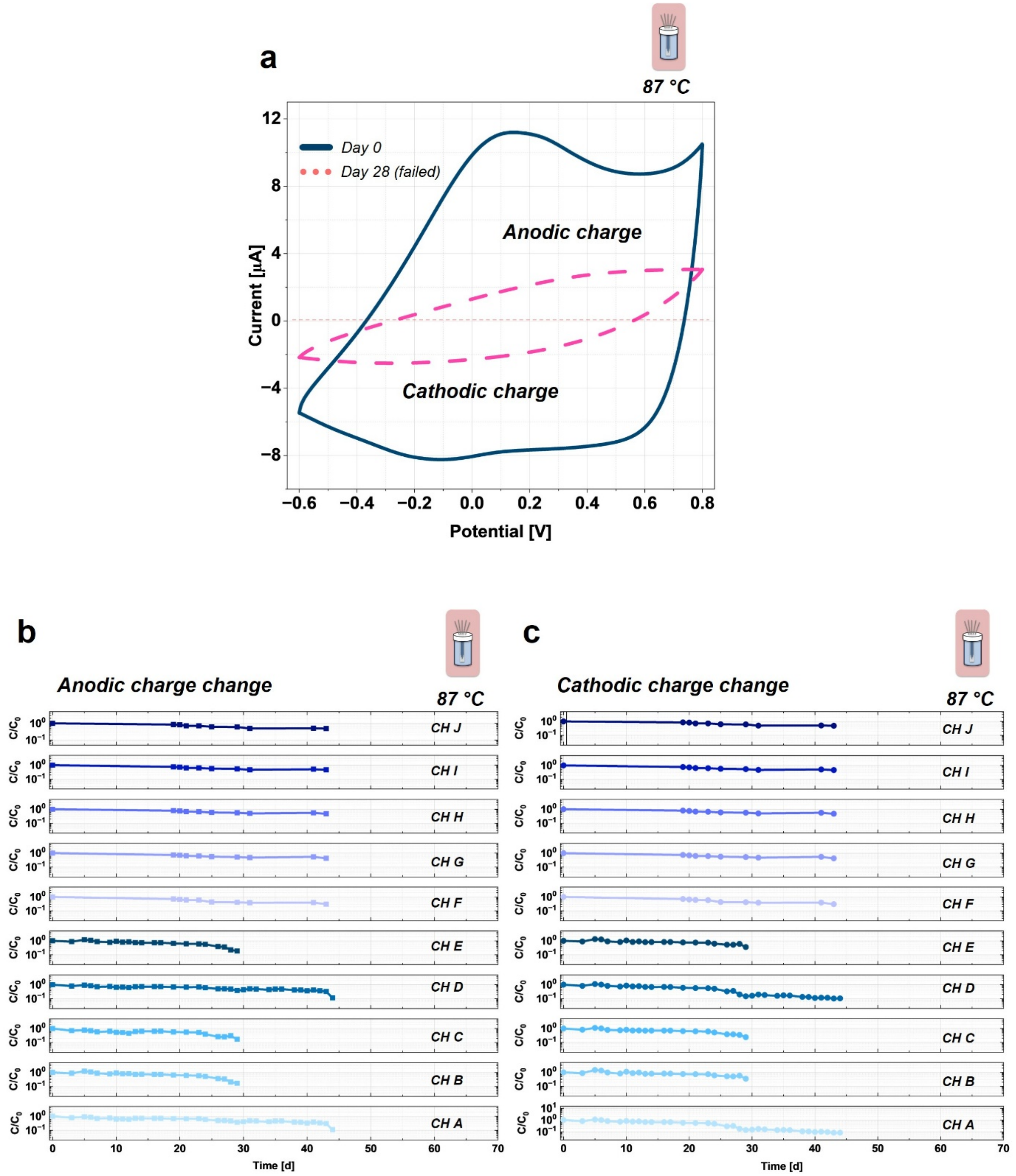


**Figure S16.** a) Representative CV curves of post-fabrication and failed devices at 87 °C. Charge change over time for 10 electrodes: b) anodic and c) cathodic charge change at 87 °C.

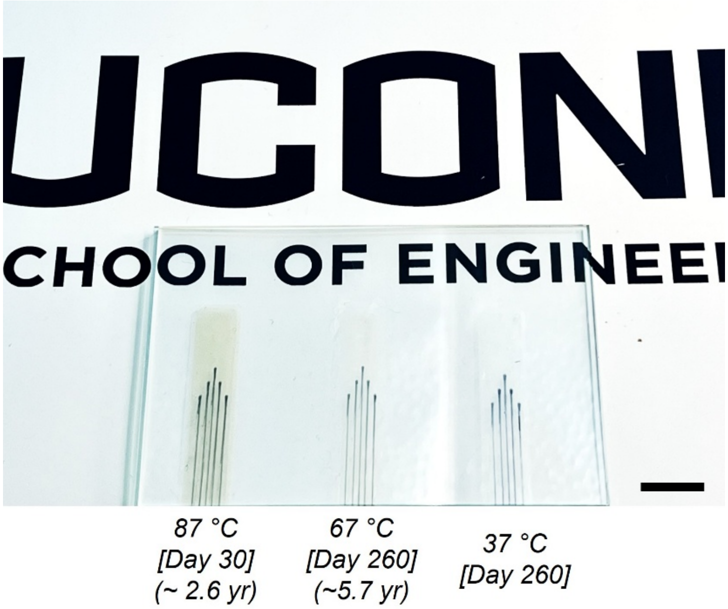


**Figure S17.** Photographs of the device under different temperature groups. Scale bar: 1 cm.

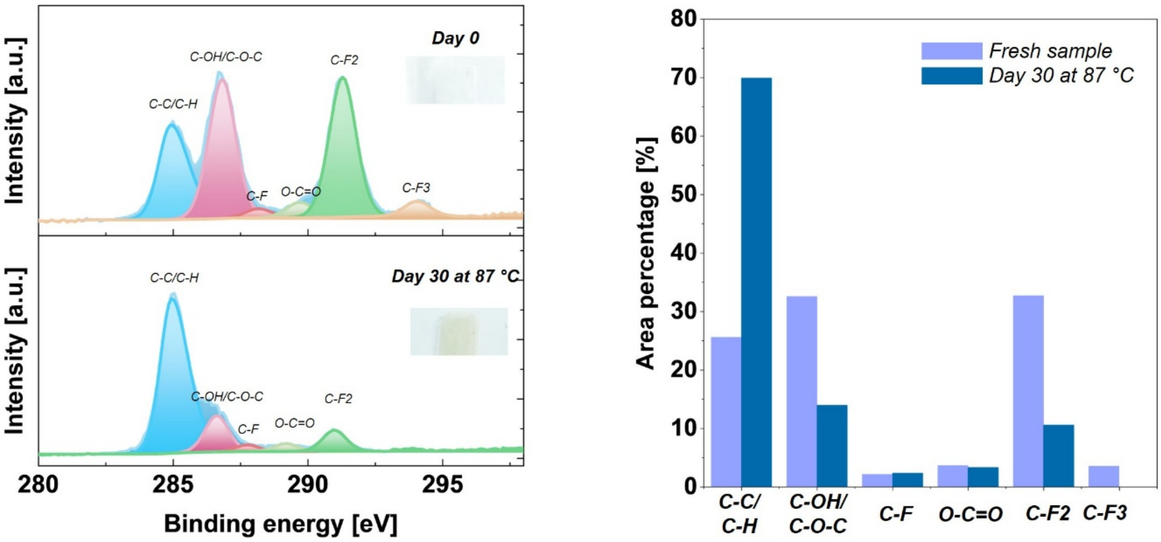


**Figure S18.** XPS analysis of fresh PVDF-HFP film and after 30 days in 1× PBS at 87 °C.

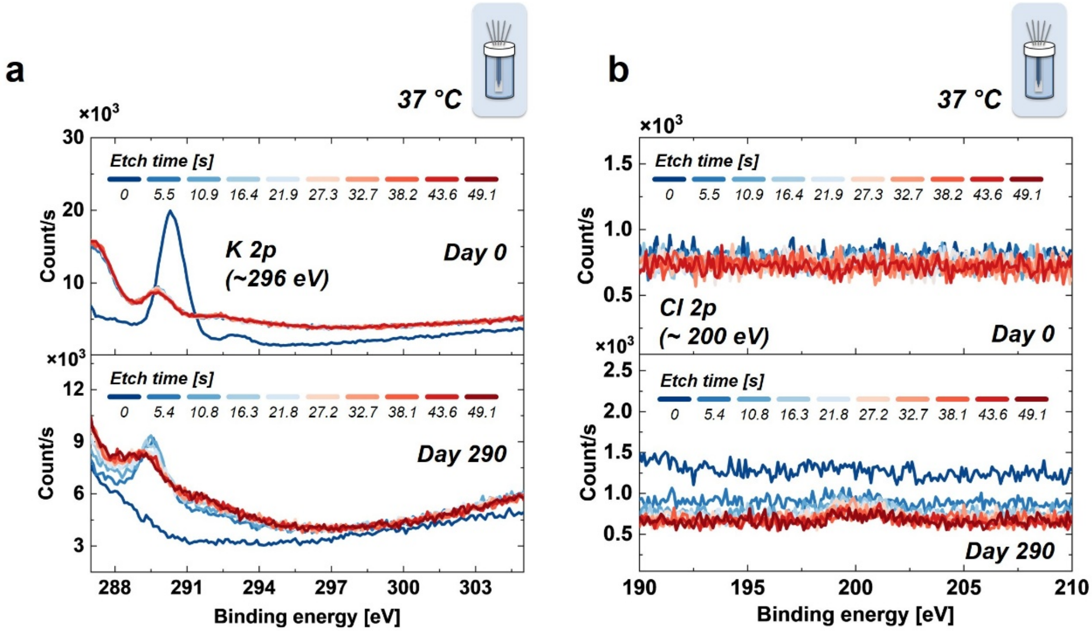


**Figure S19.** XPS analysis results before (top) and after soaking (bottom) in 1X PBS at 37 °C for 290 days: a) K 2p, b) Cl 2p.

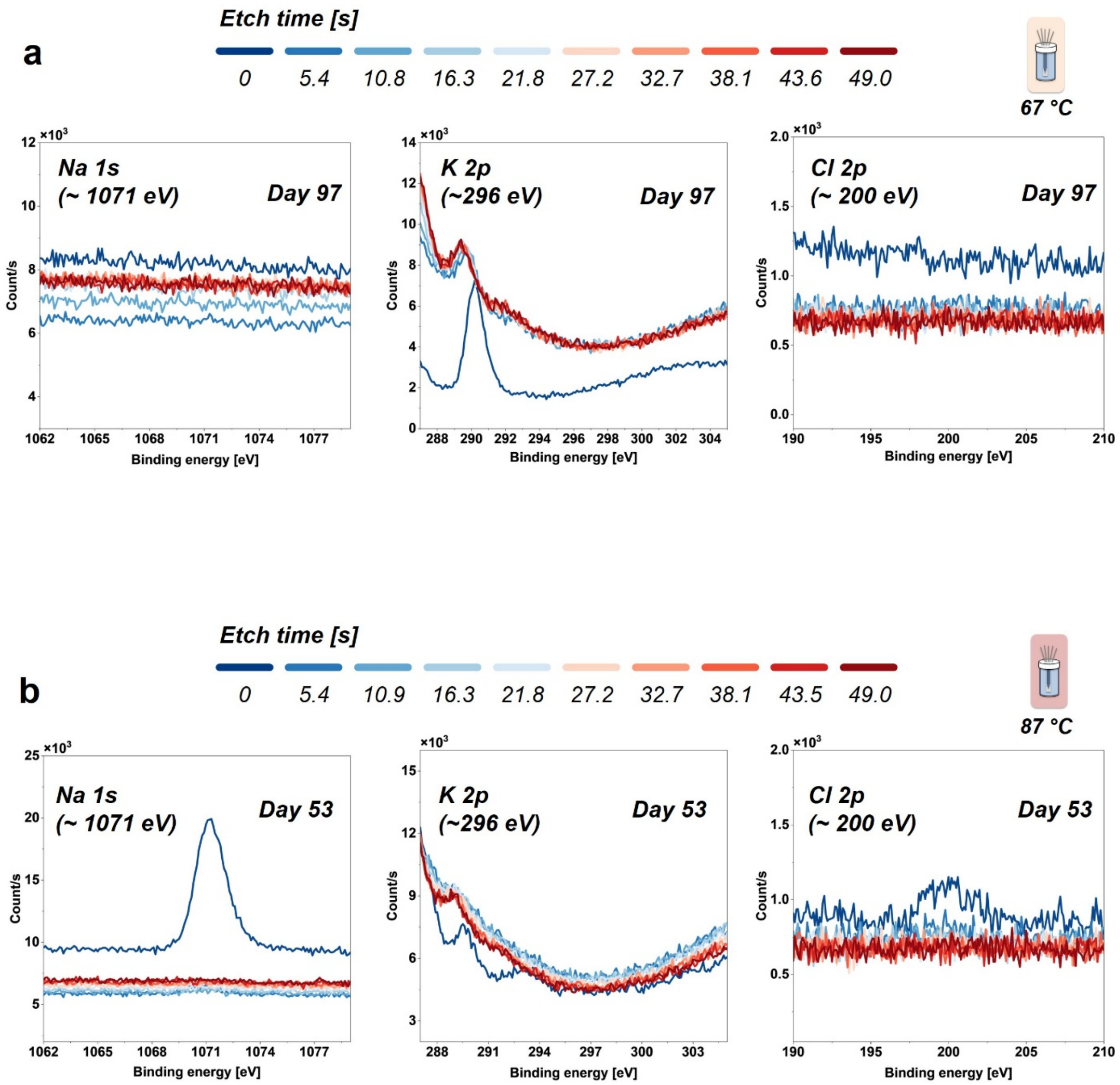


**Figure S20.** XPS spectra of Na 1s (left), K 2p (middle), and Cl 2p (right) after soaking in 1× PBS at two temperatures: a) Day 97 at 67 °C, b) Day 53 at 87 °C.

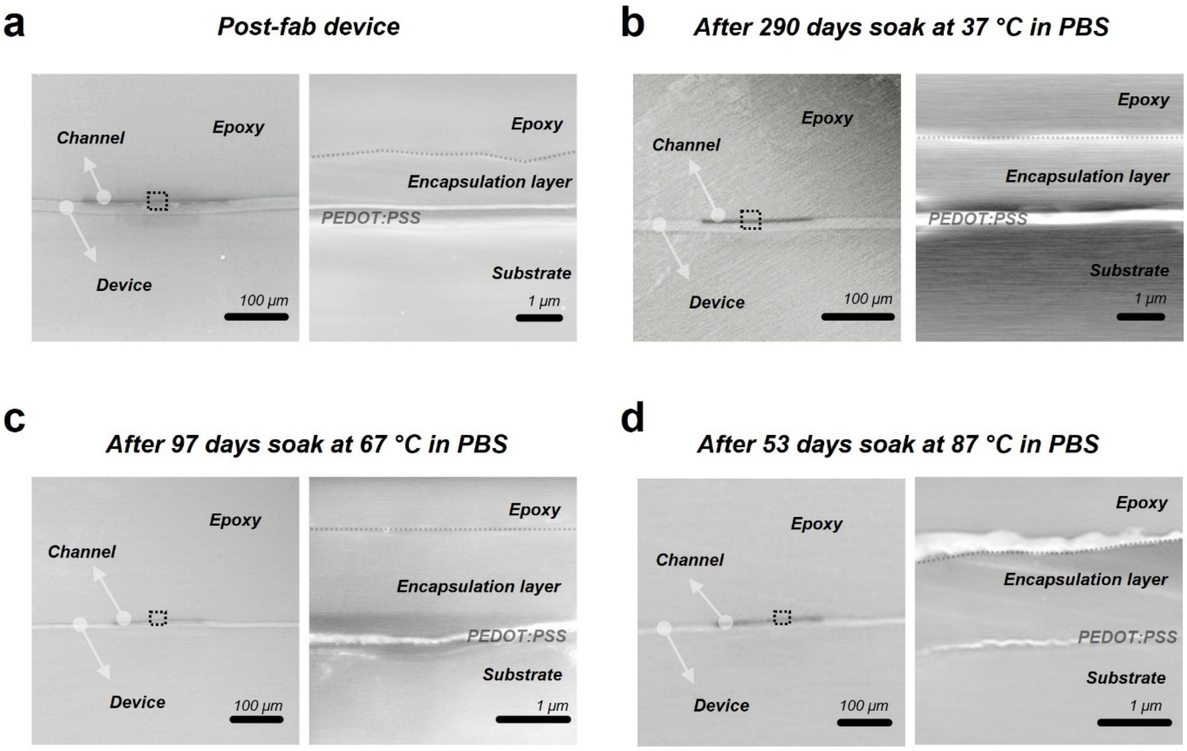


**Figure S21.** Cross section images of samples: a) Post-fabrication, b) 290 days soaked at 37 °C in PBS, c) 97 days soak at 67 °C in PBS, d) 53 days soak at 87 °C in PBS

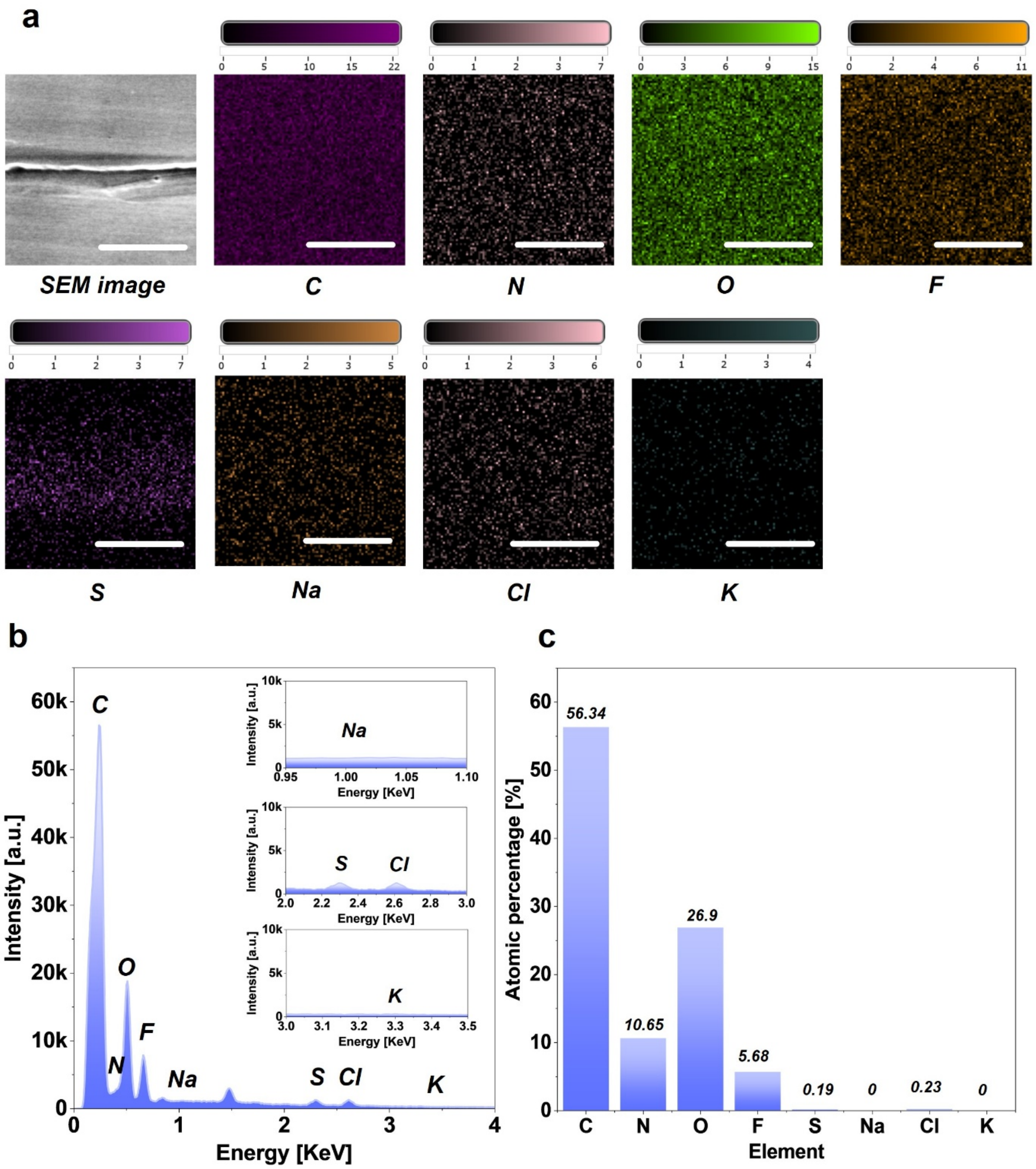


**Figure S20.** Cross section SEM—EDS analysis of post-fab device: a) Mapping results, b) SEM—EDS spectrum, c) Atomic percentage of each element. Scale bars: 5 µm.

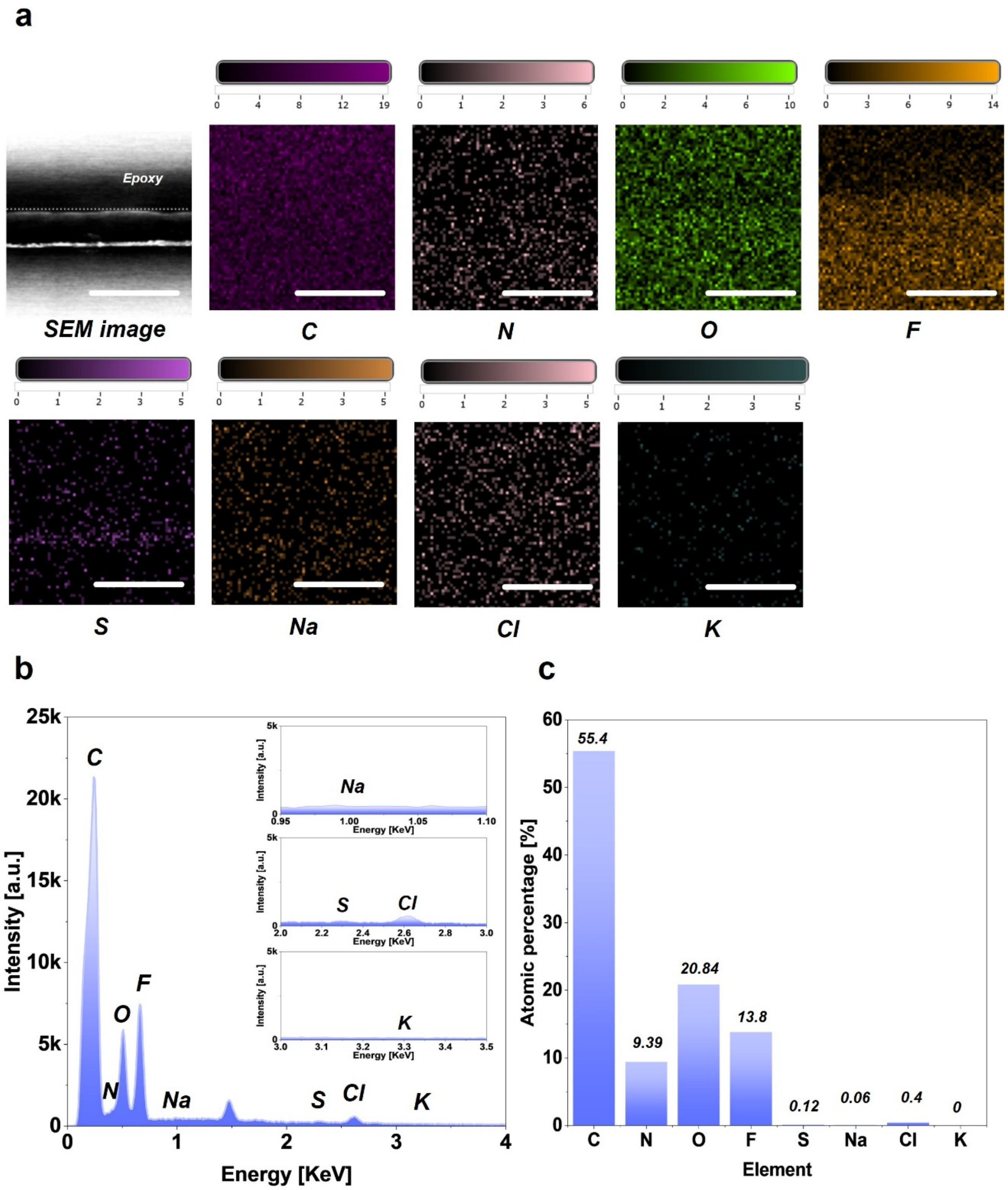


**Figure S21.** Cross section SEM—EDS analysis of device soaked in PBS for 290 days at 37 °C : a) Mapping results, b) SEM—EDS spectrum, c) Atomic percentage of each element. Scale bars: 5 µm.

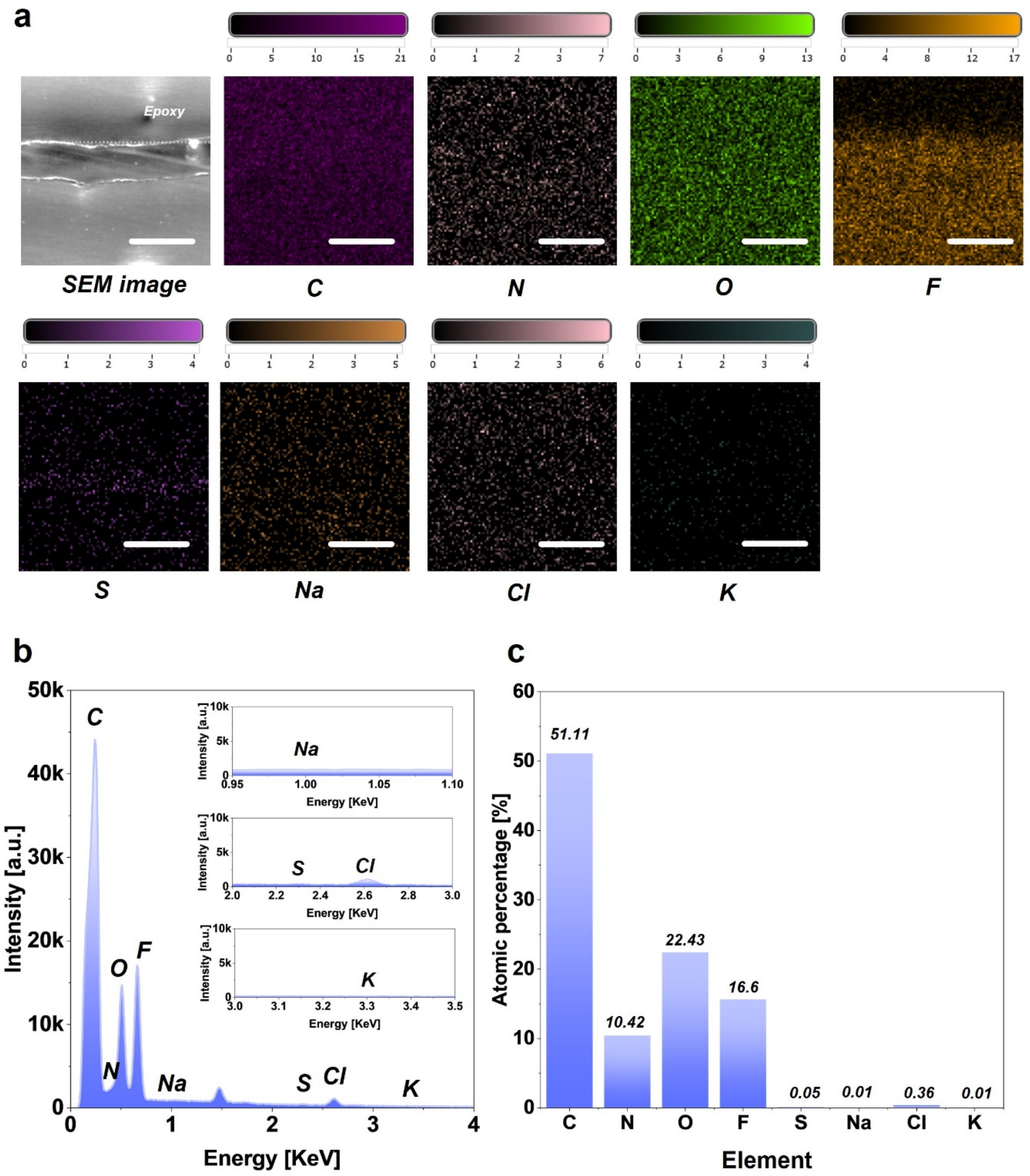


**Figure S22.** Cross section SEM—EDS analysis of device soaked in PBS for 97 days at 67 °C : a) Mapping results, b) SEM—EDS spectrum, c) Atomic percentage of each element. Scale bars: 5 µm.

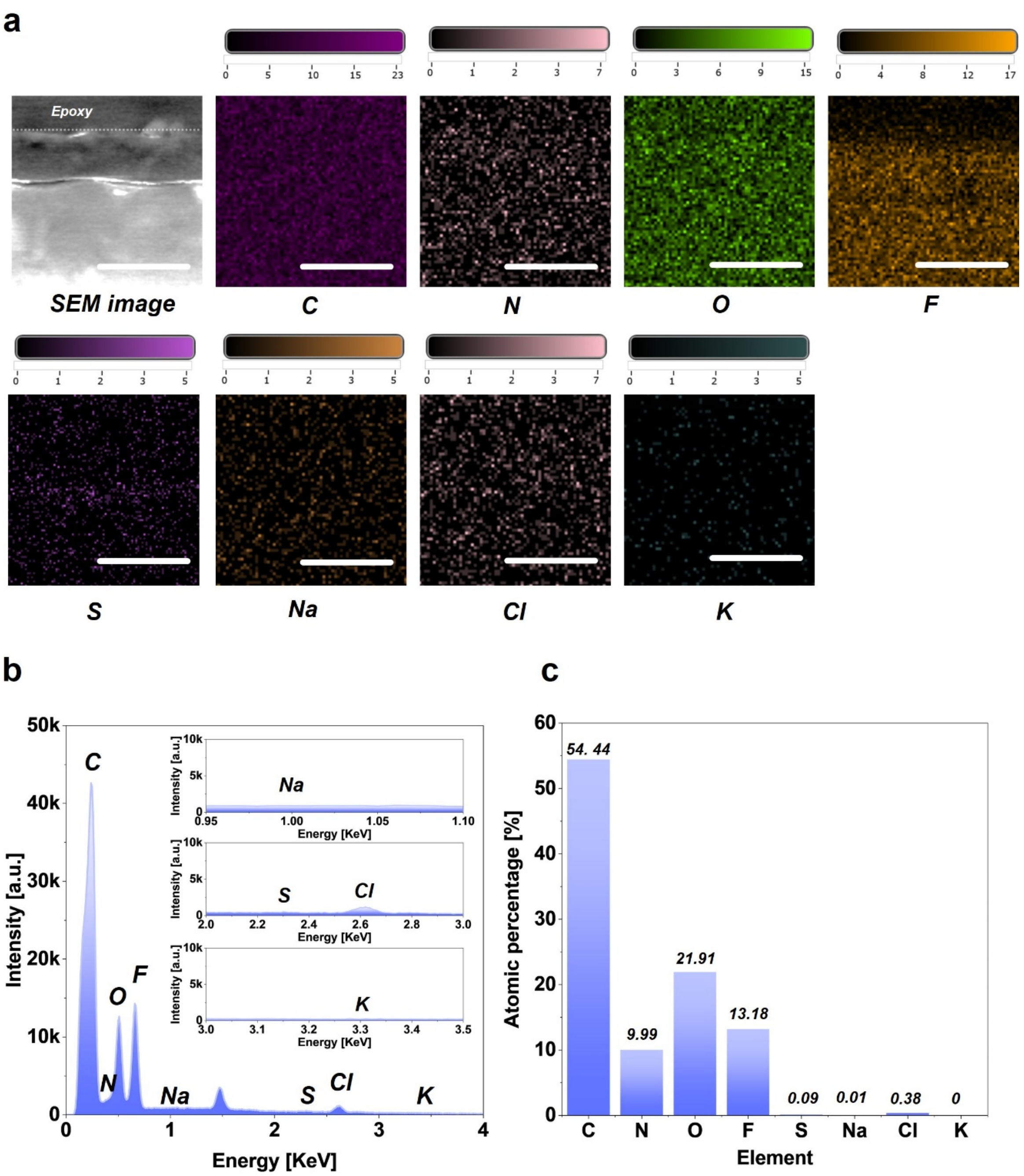


**Figure S23.** Cross section SEM—EDS analysis of device soaked in PBS for 53 days at 87 °C : a) Mapping results, b) SEM—EDS spectrum, c) Atomic percentage of each element. Scale bars: 5 µm.

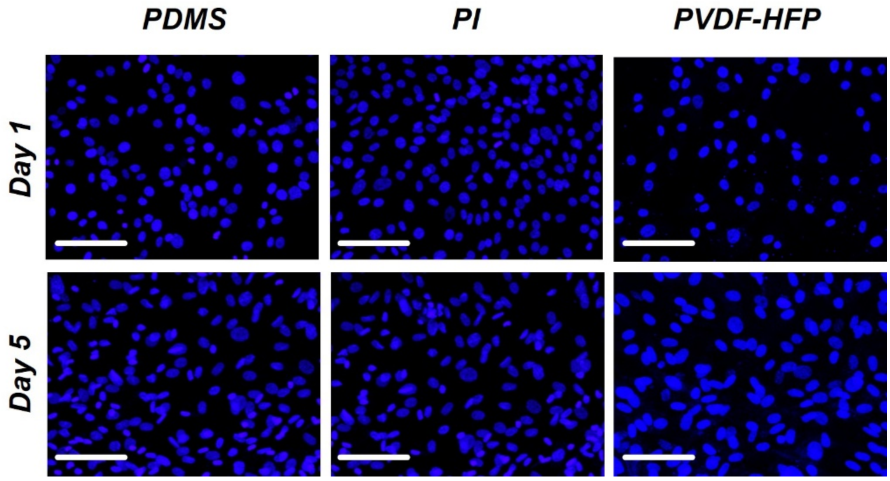


**Figure S24.** Representative confocal images of fibroblast cells stained with DAPI and cultured on three different substrate materials on Day 1 and Day 5. Scale bars: 100 µm.
